# Functional profiling reveals a non-enzymatic role of NUDT5 in repressing purine *de novo* synthesis

**DOI:** 10.1101/2025.03.16.643506

**Authors:** Jung-Ming G. Lin, Tuan-Anh Nguyen, Anne-Sophie M. C. Marques, Lorenzo Scrofani, Diana Daum, Ludwig G. Bauer, Carol Cheng, Luna D’Angelo, Juan Sanchez, Christoph Bueschl, Pisanu Buphamalai, Marton Siklos, Jakob-Wendelin Genger, Gerald Hofstaetter, Kathrin Runggatscher, Bettina Guertl, Yusi Liu, Jesper Hansen, Anna Koren, D. Sean Froese, David S. Rosenblatt, Kristaps Klavins, Andreas Bergthaler, Joerg Menche, J. Thomas Hannich, Sara Sdelci, Kilian V. M. Huber, Stefan Kubicek

## Abstract

Folate metabolism is intricately linked to purine *de novo* synthesis through the incorporation of folate-derived one-carbon units into the purine scaffold. Here, we investigate the chemical and genetic dependencies caused by mutations in the folate enzyme MTHFD1 and discover a key role for Nudix hydrolase 5 (NUDT5) in regulating purine *de novo* synthesis. Through genetic knockout and development of a selective chemical NUDT5 degrader, we uncover an unprecedented scaffolding role rather than NUDT5 enzymatic activity is responsible for this phenotype. We find that NUDT5 interacts with the rate-limiting enzyme of purine de novo synthesis, PPAT, to repress the pathway in response to elevated purine levels. Our findings establish NUDT5 as an important regulator of purine *de novo* synthesis and elucidate its role in mediating sensitivities to 6-thioguanine in cancer treatment and to adenosine in MTHFD1 deficiency.

## Main Text

Folate metabolism is essential for providing one-carbon units to the biosynthesis of numerous metabolites, including purines, thymidylate, and methionine (*1*). Consequently, impairment of folate metabolism via genetic mutations or dietary folate deficiency causes varied pathologies including developmental defects and an increased risk of cancer (*2–6*). Therapeutically, the dependence of proliferating cells on this pathway is exploited by the use of antifolates in cancer therapy (*7–9*).

The folate pathway is compartmentalized between cytoplasm and mitochondria, with additional roles of selected enzymes in the nucleus (*10, 11*). The usual direction of the folate catalytic cycle of mitochondrial formate production and cytosolic formate utilization can be reversed following mutation of key folate enzymes (*12*). Under these conditions, as well as following pharmacological inhibition with anticancer drugs like methotrexate, destabilization of the folate scaffold might further contribute to the overall phenotype (*13, 14*). Folate deficiency can also arise from disruption of downstream pathways by “trapping” of all cellular folates as certain species, e.g., as 5-methyltetrahydrofolate (5-meTHF) in vitamin B12 deficiency (*15, 16*) and as 10-formyltetrahydrofolate (10-CHO-THF) following pharmacological inhibition of MTHFD1 (*17*).

MTHFD1 (C-1 tetrahydrofolate synthase) is a trifunctional enzyme that catalyzes the cytoplasmic interconversion of 10-CHO-THF, methenyltetrahydrofolate (CH^+^-THF), and methylenetetrahydrofolate (CH_2_-THF) by its formyltetrahydrofolate synthetase (S) and methylenetetrahydrofolate dehydrogenase/methenyltetrahydrofolate cyclohydrolase domains (D/C) (Fig. 1a). While CH_2_-THF provides one-carbon units for the synthesis of methionine and thymidylate, 10-CHO-THF delivers two of the carbons to the purine scaffold generated in the *de novo* synthesis pathway. This process might be enhanced by direct interaction of MTHFD1 with the purinosome (*18*). Knockout of MTHFD1 in yeast and human cells results in purine auxotrophy, presumably by preventing purine *de novo* synthesis (*19–24*). In contrast, in cells from patients with MTHFD1 deficiency, purine *de novo* synthesis is unaffected, while thymidylate and methionine biosynthesis are impaired due to point mutations in the cyclohydrolase domain of the enzyme (*25*). To date, no comprehensive studies of the metabolic, pharmacologic, and genetic dependencies caused by loss of distinct MTHFD1 enzymatic activities have been reported. In an attempt to address this gap in our current understanding, we here show that distinct features of MTHFD1 control a switch between cellular adenosine dependency and toxicity. Importantly, we find that the antiproliferative effect of adenosine and other purine analogs in this context is dependent on a previously unidentified scaffolding function of NUDT5, revealing this NUDIX family member as a direct regulator of *de novo* purine synthesis.

**Fig. 1.**
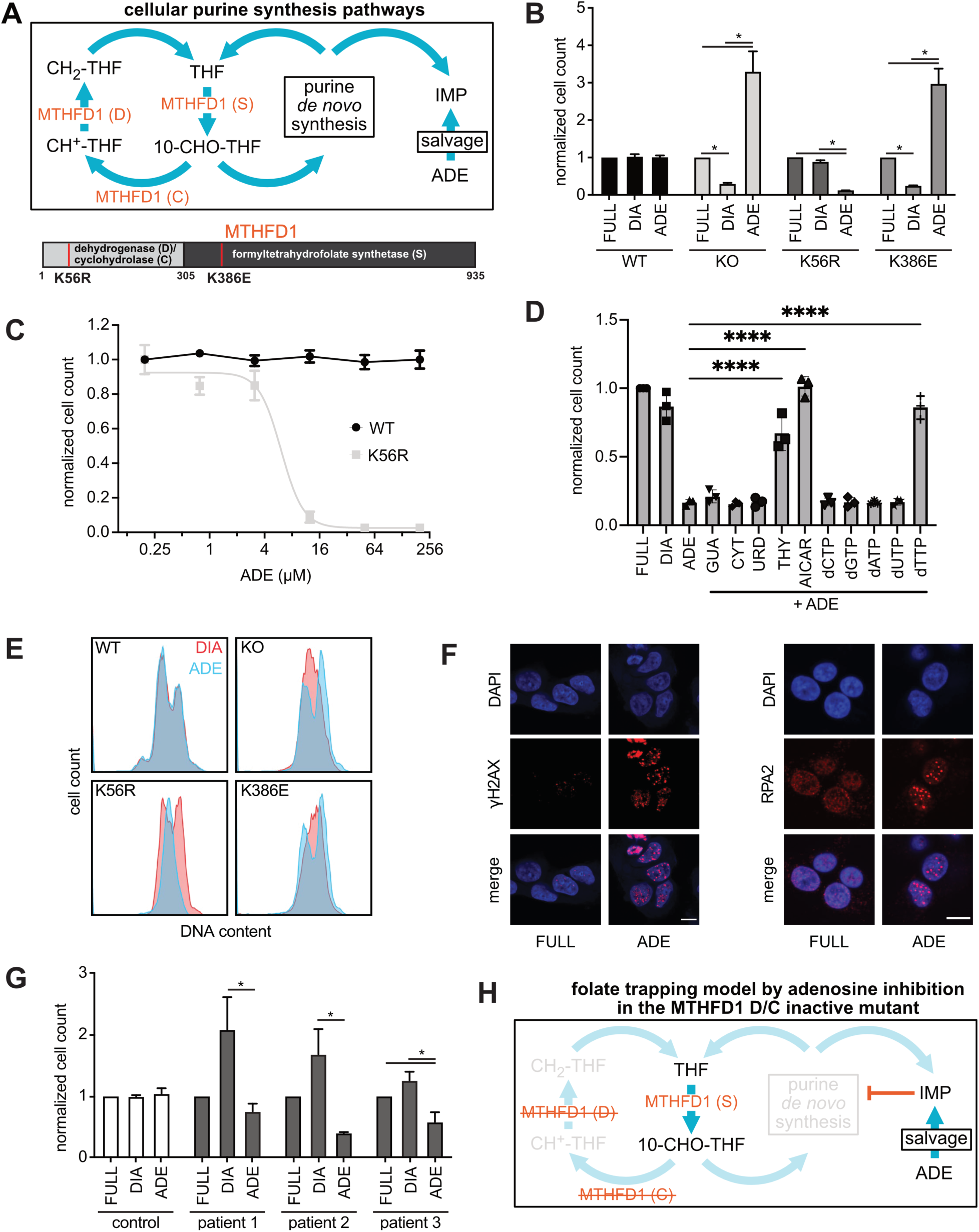
MTHFD1 enzymatic functions control a switch between adenosine dependency and toxicity. **(A)** Schematic overview of the folate cycle and cellular purine synthesis pathway. Corresponding folate intermediates are generated by the enzymatic functions of the MTHFD1 synthetase (S), cyclohydrolase (C) and dehydrogenase (D) domains. 10-CHO-THF is used for the synthesis of IMP via the purine *de novo* synthesis pathway. Alternatively, IMP can be synthesized through the salvage pathway in presence of exogenous adenosine. The schematic domain structure of MTHFD1 is shown below with sites of K56R and K386E mutations that ablate the enzymatic reactions of each respective domain. **(B)** Normalized cell counts of WT, MTHFD1^KO^, MTHFD1^386E^, and MTHFD1^K56R^ cells grown for 72 h in media containing dialyzed serum (DIA) or dialyzed serum supplemented with 50 µM adenosine (ADE), normalized to the respective clone grown in media containing full serum (FULL), (n=3 biological replicates, mean ± SD, *p<0.05). **(C)** Normalized cell counts of WT and MTHFD1^K56R^ cells at indicated concentrations of adenosine. Cells were grown for 72 h media containing dialyzed serum (DIA) or dialyzed serum supplemented with respective adenosine concentrations and normalized to the lowest adenosine concentrations. (n=2 biological replicates, mean ± SD). **(D)** Normalized cell counts of MTHFD1^K56R^ cells cultured at indicated media conditions and with supplemented nucleotides and precursors (50 µM). **(E)** Representative cell cycle plots for WT and MTHFD1 mutant HAP1 cells in DIA and ADE conditions. **(F)** Representative images of γH2A.X and RPA2 staining of MTHFD1^K56R^ cells with adenosine supplementation. Scale bar is 10 µm. **(G)** Growth assays of fibroblasts derived from MTHFD1 deficiency patients in FULL, DIA and ADE (n=3 biological replicates, Mean ± s.d, *p<0.05). **(H)** Schematic overview of the genetic folate trap model.

### Distinct MTHFD1 enzymatic activities control a switch between adenosine dependency and toxicity

We engineered wild-type HAP1 cells (WT) and derived clonal cell lines that either contain a deletion in the MTHFD1 gene (MTHFD1^KO^), or have been reconstituted with either the wild-type allele (MTHFD1^recon^) or catalytically inactive point mutations (*24, 26*) of the synthetase (MTHFD1^K386E^) or cyclohydrolase (MTHFD1^K56R^) domains, respectively (Fig. 1a). We then tested the proliferation of these cell lines in standard media conditions and in media containing dialyzed serum deprived of metabolites smaller than 10 kDa. MTHFD1^KO^ and MTHFD1^K386E^ cells were highly sensitive to serum dialysis, whereas the growth of WT, MTHFD1^recon^ and MTHFD1^K56R^ cells was unaffected in these conditions (Fig. 1b and fig. S1). To identify compounds that can counteract the detrimental effects of dialyzed serum on the growth of MTHFD1^KO^ cells, we performed a chemical screen of more than 90,000 structurally diverse small molecules (fig. S1). We realized that mainly adenine-containing compounds (e.g. NADH, NAD, FAD, SAM, AMP) or adenine precursors (e.g. hypoxanthine), but not other purine or folate analogues were able to rescue the growth of MTHFD1^KO^ cells (fig. S1). Similarly, adenosine addition reestablished proliferation of S-domain inactive MTHFD1^K386E^ cells. However, in MTHFD1^K56R^ cells harboring the inactivating mutation in the D/C domain, we observed the diametrically opposite phenotype, as in that context addition of adenosine caused a strong antiproliferative response (Fig. 1b). These effects occur in a concentration-dependent manner, where MTHFD1^K56R^ cells become already sensitized to adenosine at low micromolar concentrations (IC_50_ ∼6 µM, Fig. 1c). Moreover, the combinatorial growth deficiencies were not specific to HAP1 cells, as we could phenocopy these effects in other cancer cell lines when cells were grown in adenosine-containing media together with the MTHFD1 D/C domain inhibitor LY345899 (*26*) (fig. S2). Supplementation of thymidine, dTTP or AICAR were able to reverse these toxic effects in MTHFD1^K56R^ cells (Fig. 1d).

The addition of adenosine caused strong cell cycle changes, with MTHFD1^KO^ and MTHFD1^K386E^ cells responding with adenosine-dependent G1/S arrest to dialyzed serum conditions, whereas MTHFD1^K56R^ cells arrest in G1/S following adenosine addition (Fig. 1e, fig. S3a). Drastic increases in foci formation for both RPA and γH2A.X in these conditions (Fig. 1f, fig. S3b) suggest that replication stress and DNA damage cause this phenotype. We further note that these effects occur at adenosine concentrations that are orders of magnitude lower than the millimolar concentrations normally needed to induce replication stress through nucleotide imbalance (*27*). Again, this was not specific to HAP1 cells, as we observed similar DNA damage effects in fibroblasts from MTHFD1-deficient patients (Fig. 1g, fig. S4).

Overall, by engineering a specific mutation that inactivates the D/C domain of MTHFD1, we have established a genetic “folate trapping” model (*17*) (i.e. accumulation of 10-CHO-THF), where presumably regeneration of THF and thus maintenance of balanced folate and nucleotide levels are prevented by the concomitant inhibition of the folate cycle (through MTHFD1^K56R^ mutation) and of purine *de novo* synthesis (by adenosine addition) (Fig. 1h, fig. S5), thereby leading to DNA damage and ultimately cell death. We hypothesized that this phenotype could be rescued by genetic and chemical modulators of the folate and purine *de novo* synthesis pathways.

### A genome-wide genetic screen identifies NUDT5 as modulator of MTHFD1-mediated adenosine response

We performed a genome-wide genetic loss-of-function screen to probe genetic dependencies of the folate-trap conditions. For this, we transduced MTHFD1^K56R^ cells with the Brunello knockout library targeting 19,114 human genes with an average of four sgRNAs per gene. Following selection and growth with an, in that context, toxic adenosine concentration of 50 μM for two weeks, we analyzed sgRNA abundance as a proxy for beneficial growth effects following knockout of the respective target gene (Fig. 2a).

**Fig. 2.**
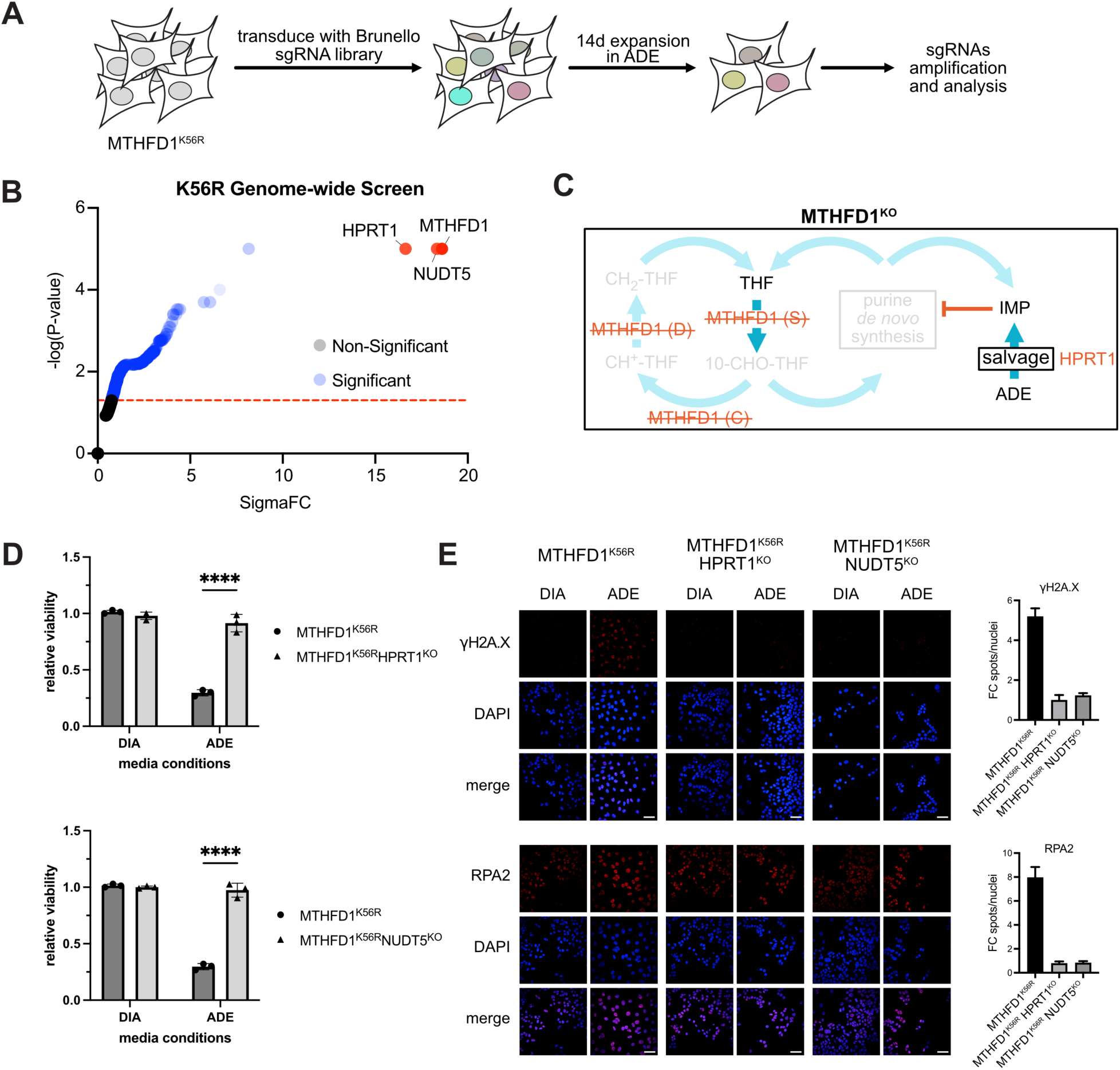
A genome-wide knockout screen identifies folate trap modulators. **(A)** Schematic overview of the genetic screening strategy. **(B)** Results of the genome-wide KO screen as performed in a), highlighting significantly depleted genes (HPRT1, NUDT5, MTHFD1). **(C)** Model of folate trap rescue by MTHFD1 knocked-out in MTHFD1^K56R^ cells. **(D)** Normalized viability of MTHFD1^K56R^HPRT1^KO^ and MTHFD1^K56R^NUDT5^KO^ cells cultured in media containing dialyzed serum (DIA) or the same media supplemented with 50 µM adenosine (ADE). Viability was measured after 72 h and normalized to respective DIA conditions. (n=2, mean ± s.d., two-way ANOVA, ****p<0.0001). **(E)** Representative images of γH2A.X and RPA2 staining of MTHFD1^K56R^, MTHFD1^K56R^HPRT1^KO^ and MTHFD1^K56R^NUDT5^KO^ cells after being cultured in DIA or ADE for 72 h. Spots/nuclei values were normalized to DIA conditions. Scale bar is 50 µm.

We discovered that the knockout of three genes was able to rescue adenosine-mediated toxicity: MTHFD1, HPRT1 (hypoxanthine phosphoribosyl transferase 1), and NUDT5 (Fig. 2b). MTHFD1 served as a positive control, as a shift from the MTHFD1^K56R^ mutation to the full knockout was expected to correlate with the transition from adenosine toxicity to adenosine dependency (Fig. 1b and 2c). HPRT1 is an enzyme in the purine salvage pathway that plays a key role in recycling purines (*28, 29*). We validated the absence of purine-mediated toxicities in MTHFD1^K56R^HPRT1^KO^ double-mutant cells and showed that in these cells adenosine does not cause DNA damage signals (Fig. 2d and e, fig. S6a and b). These data indicate that the functional utilization of exogenous adenosine in intracellular biochemical pathways is required to trigger the folate trap, consistent with the model that cellular modulation of the purine salvage pathway plays a key role in repressing purine *de novo* synthesis activity (Fig. 2d and e, fig. S6b).

Thus, we turned our attention to the third hit NUDT5 and confirmed that its knockout in the MTHFD1^K56R^ background (MTHFD1^K56R^NUDT5^KO^) reestablished cellular proliferation in purine-rich media (Fig. 2d and e, fig. S6a and b). NUDT5 is a hydrolase involved in the catabolism of ADP-ribose and nuclear ATP synthesis (*30–34*), but so far it has not been directly linked to purine sensing or folate metabolism (fig. S6b).

### NUDT5 is required for the repression of de novo purine biosynthesis in response to adenosine

To assess possible NUDT5 mechanisms of action, we first performed comparative metabolomics in WT, MTHFD1^K56R^, NUDT5^KO^ and MTHFD1^K56R^NUDT5^KO^ cells. We used isotope tracing to monitor whether selected metabolites originate from cellular purine salvage or *de novo* synthesis activity, respectively. Specifically, we treated cells with isotope-labeled formate (+1 ^13^C) that is incorporated into *de novo* synthesized purines via 10-CHO-THF produced by the MTHFD1 synthetase domain (Fig. 3a). To assess incorporation of externally added adenosine into different metabolites, we used isotope labeled adenosine (+1 ^15^N). Performing metabolome analysis by high resolution mass spectrometry enabled us to clearly discriminate metabolites derived from *de novo* synthesis (containing heavy carbon atoms) from those derived from adenosine salvage (containing heavy nitrogens).

**Fig. 3.**
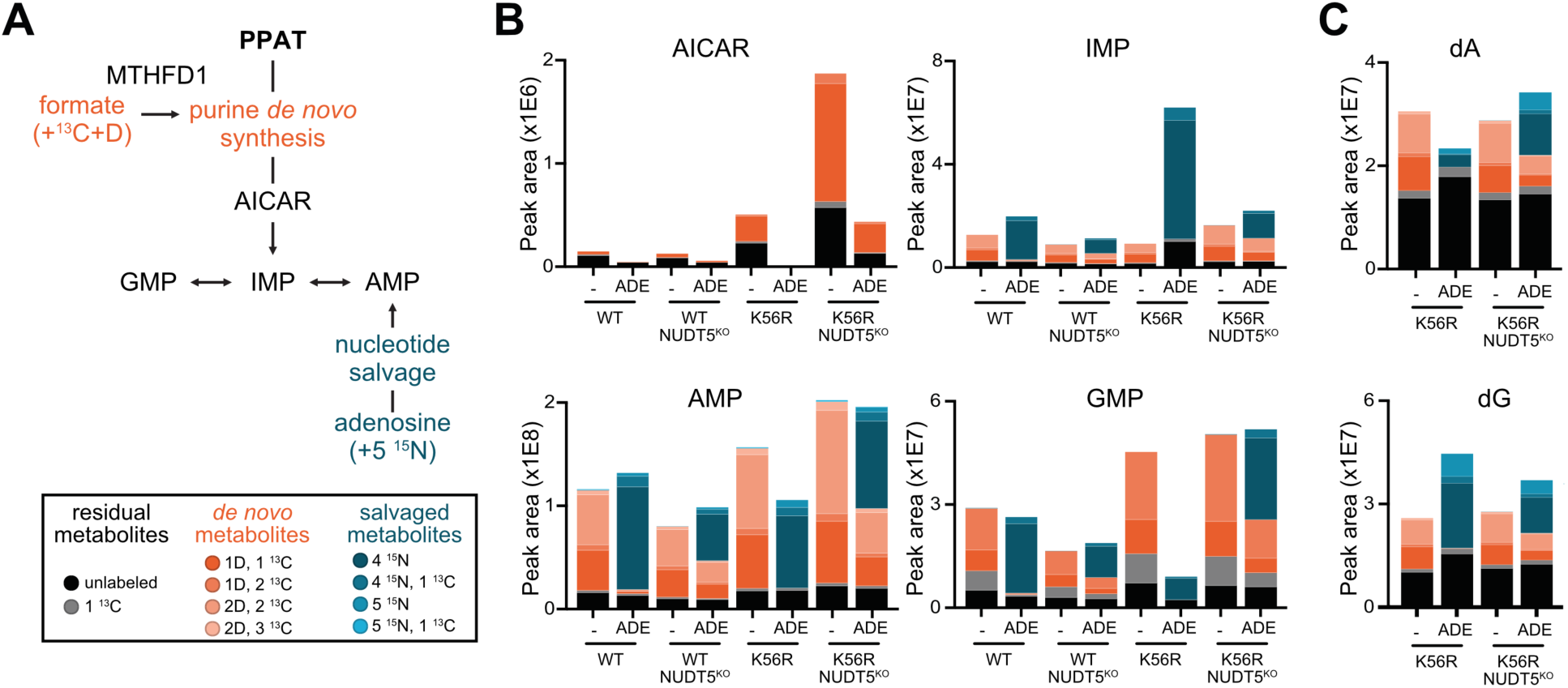
Loss of NUDT5 causes increased *de novo* purine synthesis in the presence of salvageable adenosine. **(A)** Schematic overview of the purine *de novo* synthesis and salvage pathways indicating labeled precursors used in isotope tracing experiments. WT, NUDT5^KO^, MTHFD1^K56R^ and MTHFD1^K56R^NUDT5^KO^ cells were grown in DIA containing 1 mM labeled formate, optionally supplemented with 50µM labeled adenosine (ADE) for 24 h. **(B)** Peak areas for metabolites AICAR, IMP, AMP and GMP, labeled by detected isotope indicating metabolites derived from purine *de novo* synthesis in shades of red and those derived from salvaged adenosine in shades of blue. **(C)** Peak areas corresponding to dA and dG obtained from hydrolyzed DNA, colored as in panel b.

For all measured metabolites (IMP, AMP, GMP), adenosine addition resulted in a strong repression of *de novo* synthesis in both WT and MTHFD1^K56R^ cells (Fig. 3b). Knock-out of NUDT5 prevented this repression, both in WT and in MTHFD1^K56R^ cells, and resulted in a drastically elevated proportion of *de novo*-derived AMP, GMP and IMP (Fig. 3b). The same effect was also clearly observable in nucleotides derived from hydrolyzed DNA, where the lower turnover of DNA resulted in a higher fraction of unlabeled nucleotides (Fig. 3c). Together, these data indicate that loss of NUDT5 – irrespective of MTHFD1 mutational status – enables cellular purine production through the *de novo* pathway even when exogenous purines are abundant.

### A NUDT5 scaffolding function rather than enzymatic activity is essential for modulating adenosine responses

Because NUDT5 can be broadly linked to purine metabolism throughout its ADP-ribose converting activity (*30, 32*), we next investigated whether its enzymatic activity plays a central role in *de novo* purine synthesis repression. For this, we first performed viability assays using the well-characterized chemical inhibitor (*35*) TH5427 in MTHFD1^K56R^ cells. To our surprise, we found that even at doses more than 1,000x higher than its biochemical IC_50_ of 29 nM, TH5427 treatment did not rescue the viability of MTHFD1^K56R^ cells when cultured in high adenosine-containing media (Fig. 4a).

**Fig. 4.**
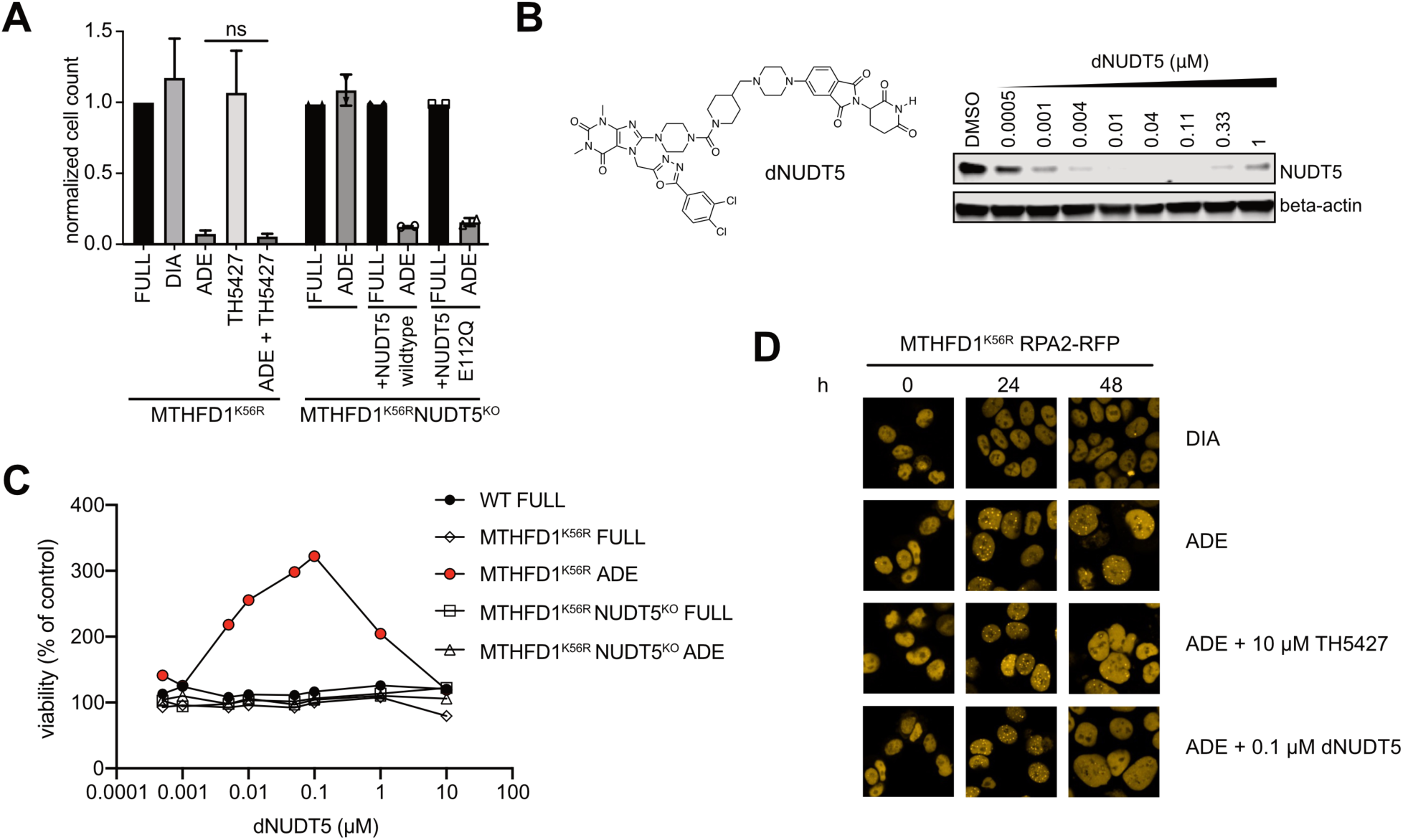
Knockout or chemical degradation, but not enzymatic inhibition, of NUDT5 prevents adenosine-mediated toxicity. **(A)** Normalized cell counts of MTHFD1^K56R^ and MTHFD1^K56R^ NUDT5^KO^ cells treated for 72 h in ADE and 10 µM NUDT5 inhibitor TH5427 alone and in combination. **(B)** Chemical structure of the NUDT5 targeting PROTAC dNUDT5. Representative Western blot of dose-dependent dNUDT5 effects on NUDT5 protein levels after 20 h treatment in WT cells. **(C)** Dose-dependent effects of dNUDT5 on the growth of WT and MTHFD1^K56R^ cells cultivated in FULL or DIA for 72 h. **(D)** Representative images of RPA2-foci formation of MTHFD1^K56R^ RPA2-RFP (intron tagged) cells at indicated media conditions and times.

We then performed proliferation assays in MTHFD1^K56R^NUDT5^KO^ cells reconstituted with either NUDT5^wt^ or the catalytically inactive NUDT5^E112Q^ variant (*34*). Both protein variants resensitized MTHFD1^K56R^NUDT5^KO^ cells to adenosine (Fig. 4a), corroborating our findings with the pharmacological NUDT5 inhibitor. Thus, the combined data strongly suggest that the enzymatic function of NUDT5 is dispensable for adenosine-mediated toxicity.

Reconsidering our genetic screen where the entire NUDT5 protein was lost, we speculated that the physical presence of the NUDT5 protein scaffold rather than its enzymatic activity might dictate the adenosine mediated toxicity. To dynamically probe a potential scaffolding role, we developed the small molecule tool compound dNUDT5, a PROTAC that induces targeted degradation of NUDT5 (detailed characterization in the accompanying paper submitted to Nature Chemical Biology). We confirmed that dNUDT5 induced selective NUDT5 degradation in a dose-dependent manner with a hook effect at higher concentrations that is commonly observed with PROTACs (Fig. 4b). We then performed viability assays with WT and MTHFD1^K56R^ cells in the folate trap conditions and in presence of dNUDT5 (Fig. 4c). While the compound did not affect the viability of WT cells, it prevented adenosine toxicity in MTHFD1^K56R^ through loss of accumulating DNA damage signals (Fig. 4d) and with a hook-shaped dose-responsiveness perfectly matching NUDT5 degradation. The maximum rescue of cell viability occurred at 100 nM, the same concentration that also caused maximum degradation (Fig. 4c). Altogether, we were able to phenocopy the genetic ablation of NUDT5 using an acute chemical degrader rather than an inhibitor of the enzymatic activity. The combined chemical and genetic evidence strongly suggest that the NUDT5 scaffold rather than the protein’s enzymatic activity is responsible for repressing purine *de novo* synthesis when adenosine is abundant.

### The direct interaction of NUDT5 with PPAT represses purine de novo synthesis

We next sought to identify interaction partners of NUDT5 aiming to mechanistically understand how the scaffold represses the purine *de novo* synthesis pathway.

We performed interaction proteomics experiments (*36*) with BFP-tagged NUDT5, and observed strong enrichment of three interactors: **pyrroline-5-carboxylate reductase** 1 and 2 (PYCR1, PYCR2) and phosphoribosyl pyrophosphate amidotransferase (PPAT) (Fig. 5a). The same factors were also enriched when performing NUDT5 TurboID, in chemical proteomics with a NUDT5 affinity probe (fig. S7a and b) and in unbiased large-scale protein-protein interaction data sets (*37*) performed in other cell lines. Of these interactors, we focused our attention on PPAT, since it catalyzes the rate-limiting step of the purine *de novo* synthesis pathway. We confirmed that the interaction is reciprocal, and immunoprecipitation of endogenous PPAT resulted in enrichment of NUDT5 (fig. S7c). Importantly, adenosine addition increased the strength of the NUDT5/PPAT interaction (Fig. 5b).

**Fig. 5.**
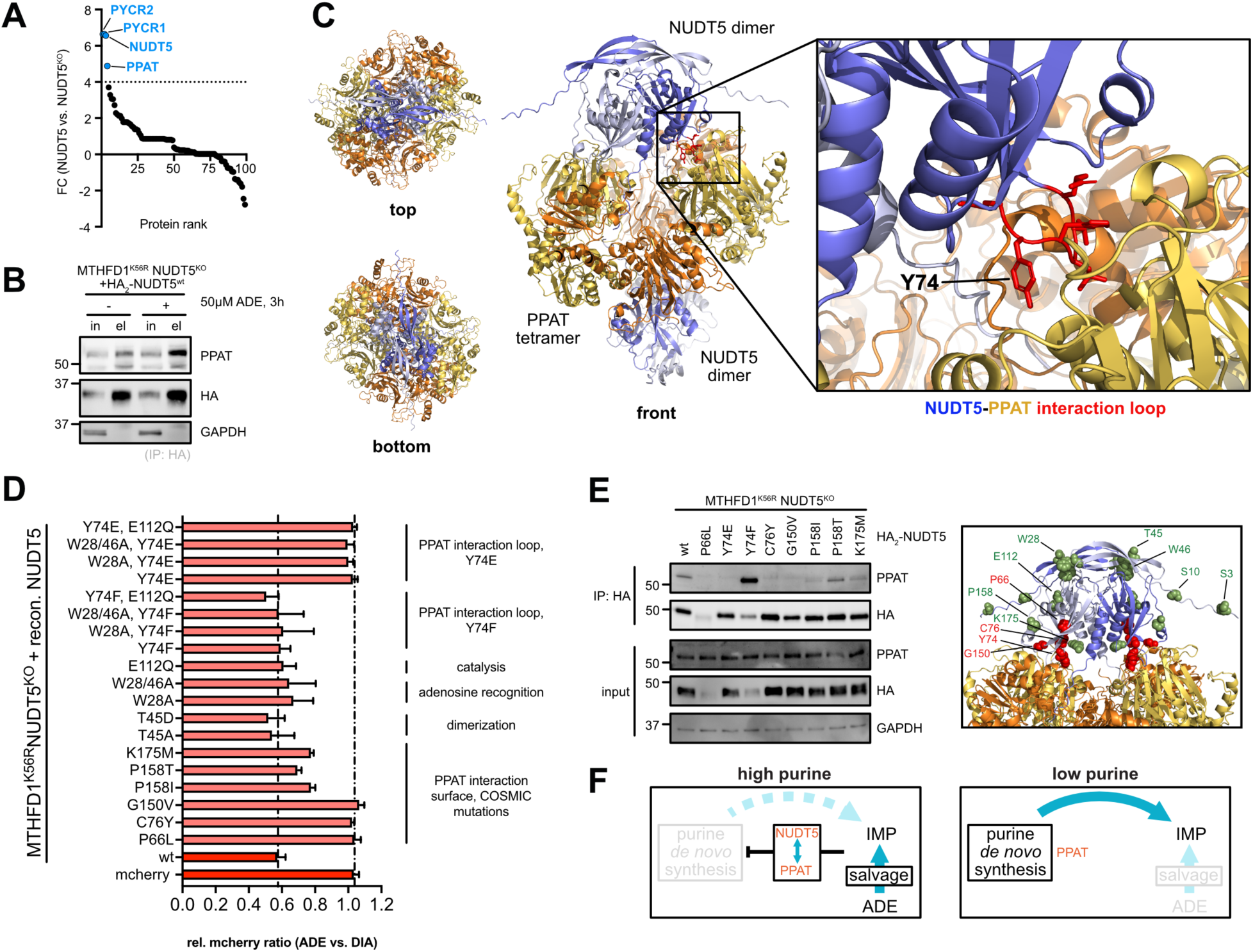
Adenosine mediated toxicity is mediated by the interaction of NUDT5 with PPAT. **(A)** Results of BFP-pulldown in MTHFD1^K56R^NUDT5^KO^ cells reconstituted with BFP-NUDT5. Hit proteins were ranked according to fold change and hits with FC >4 are highlighted in blue. **(B)** HA-Pulldown of MTHFD1^K56R^NUDT5^KO^ cells transfected with HA_2_-NUDT5^wt^ (representative blot from two independent experiments). Cells were cultivated in DIA or in ADE for 3 h. **(C)** Alphafold3 model of the NUDT5/PPAT complex with interaction site depicted in detail. **(D)** Results of fluorescence competition assay to test the effects of NUDT5 mutations on adenosine mediated toxicity in MTHFD1^K56R^ cells. Flow cytometry analysis was performed after cells were cultivated in DIA (control) or ADE for 72 h. The ratio of mcherry expressing cells was normalized to respective control conditions. Dashed lines indicate NUDT5^wt^ and mcherry control values (n=2 biological experiments, mean ± s.d.) **(E)** Pulldown of MTHFD1^K56R^NUDT5^KO^ cells transfected with respective HA-NUDT5 variants (representative blot from two independent experiments). Selected positions are depicted on the right structure on one NUDT5 subunit (Alphafold3 model). Green indicates PPAT binding permissive mutations, while mutations of the red residues resulted in loss of PPAT binding. **(F)** Proposed model of the NUDT5/PPAT dependent *de novo* synthesis pathway regulation at high and low purine conditions, respectively.

To gain initial insights into the molecular details of the NUDT5-PPAT interaction, we performed structural analyses using AlphaFold3 (*38*). These results suggest a ‘sandwich’ binding mode where two NUDT5 dimers cross-interact with a PPAT tetramer, thereby potentially locking it in an inactive conformation (Fig. 5c). We identified NUDT5^70-75^ as the putative PPAT interaction loop, within which NUDT5^Y74^ points towards the PPAT interface and might function as a key residue critical for the interaction (Fig 5c).

To test the importance of these residues, we systematically reconstituted MTHFD1^K56R^NUDT5^KO^ cells with mcherry-P2A-NUDT5 constructs carrying selected point mutations. We then performed comparative fluorescent competition assays under folate trap and control conditions (Fig. 5d and fig. S8a).

Comparable to the reintroduction of NUDT5^wt^, inactivating the enzymatic capability of NUDT5 either by abolishing recognition of the adenosine moiety (*34*) (W28A and W28/46A) or catalysis (E112Q) did not rescue the adenosine toxicity (Fig. 5d). These data corroborate our previous findings that the NUDT5 scaffold rather than the enzymatic activity is the key regulator of the phenotype. Similarly, altering the NUDT5 posttranslational modification and dimerization states (*30*) in NUDT5^T45A^ and NUDT5^T45D^ mutants as well as introduction of a NUDT5^Y74F^ mutation did not rescue the adenosine-mediated toxicity (Fig. 5d and fig. S8a and b). However, MTHFD1^K56R^ cells reconstituted with the NUDT5^Y74E^ variant were desensitized to the toxic adenosine conditions and behaved like NUDT5^KO^ cells (Fig. 5d and fig. S8b). Notably, in contrast to NUDT5^wt^ and NUDT5^Y74F^, this mutation resulted in loss of the interaction with PPAT, suggesting that the rescue of adenosine-mediated toxicity acts through release of PPAT binding (Fig. 5e).

### Loss of NUDT5-PPAT interaction confers to 6-thioguanine resistance

Antimetabolites such as purine analogs inhibit key metabolic pathways and thus are widely used in cancer treatment (*39*). We note that knock-out of HPRT1 and NUDT5 have previously been reported to confer resistance towards the clinically approved anticancer drug 6-thioguanine (6-TG) (*40*). While conceivably the knock-out of HPRT1 prevents the conversion of 6-TG to toxic downstream analogs via the purine salvage pathway, rescue of toxicity by NUDT5^KO^ has been attributed to its enzymatic function and loss of ribose-5-phosphate synthesis, which is an important building block for purine synthesis (*40*). However, we find that alike adenosine toxicity in the folate trap condition, 6-TG toxicity is similarly dependent on the presence of the NUDT5 scaffold rather than its enzymatic activity (accompanying manuscript submitted to Nature Chemical Biology and fig. S8d). These findings prompted us to explore whether NUDT5 mutations might contribute to 6-TG resistance in cancer patients. Querying the COSMIC database (*41*) for somatic NUDT5 mutations, we selected reported NUDT5 patient mutations surrounding the PPAT interaction loop and tested their influence on PPAT binding (Fig. 5e). Performing similar assays as described above, we overall observed loss of binding for all selected mutations, which either are directly facing towards the PPAT interface (C76Y, G150V) or had a destabilizing effect on NUDT5 (P66L), while the remaining mutations (P158I/T, K175M) still retained some residual PPAT binding (Fig. 5e). The binding strength was correlated with the degree of rescue of adenosine-mediated toxicity in MTHFD1^K56R^ cells (Fig. 5d and fig. S8b) and of 6-TG toxicity in HAP1^wt^ cells (fig. S8d). Thus, these data suggest that NUDT5 mutations that impair the interaction with PPAT likely contribute to antimetabolite resistance in cancer cells.

## Discussion

By initially characterizing the pharmacological and genetic dependencies caused by MTHFD1 mutations, we validated that MTHFD1^K56R^ cells constitute a genetic folate trap model in which subphysiological levels of adenosine cause strong toxicity. Utilizing this model allowed us to systematically identify modulators of the purine *de novo* synthesis pathways, leading to the discovery of a novel regulatory role for the NUDIX hydrolase NUDT5 in this context.

The folate trap model postulates that inhibition of the D/C domain of MTHFD1 results in all cellular folates accumulating as 10-CHO-THF when this metabolite is not utilized in purine *de novo* synthesis (*17*). Under these conditions, depletion of other folates, particularly CH_2_-THF that is required for thymidylate synthesis, triggers nucleotide imbalance and DNA damage. Consistently, thymidine and dTTP supplementation are able to rescue adenosine toxicity in MTHFD1^K56R^ cells. Interestingly, AICAR, which is also able to rescue cell death, is the substrate of the ATIC-mediated second 10-CHO-THF utilizing step in purine *de novo* synthesis. This suggests that the adenosine toxicity is mediated by a block of purine *de novo* synthesis before the first 10-CHO-THF utilizing step, which is mediated by GART, the enzyme directly downstream of PPAT.

The genome-wide genetic screen should therefore yield modulators of THF utilization, nucleotide salvage and early steps in *de novo* synthesis, and with MTHFD1, HPRT1 and NUDT5 we find examples for each of these functionalities. While MTHFD1 and HPRT1 could easily be functionally linked to folate metabolism and nucleotide salvage, it was not immediately evident how NUDT5 specifically affected the phenotype. Intriguingly, we found that the genetic deletion of NUDT5 enabled purine *de novo* synthesis and thus 10-CHO-THF utilization even in the presence of high adenosine concentrations, irrespective of the presence of folate trapping conditions or MTHFD1 mutations.

Importantly, we uncover that a scaffolding rather than an enzymatic role of NUDT5 is essential for repressing purine *de novo* synthesis. Extensive proteomic profiling revealed that the regulation of this pathway occurs by a direct NUDT5-PPAT interaction, with Alphafold modeling suggesting a tight interaction between two NUDT5 dimers and one PPAT tetramer. Unfortunately, we have not yet been able to experimentally obtain a co-structure of the two proteins. In this regard, it is worth noting that so far, no experimental structure for human PPAT has been reported. Likely, oxidation sensitivity of PPAT as an iron-sulfur cluster protein (*42*) contributes to the challenges in structurally resolving the complex. To validate the Alphafold model, we therefore turned to mutagenesis studies and discovered that single point mutations in the predicted interaction surface are sufficient to prevent NUDT5 binding to PPAT (Fig. 5e).

That stabilization of higher order complexes presumably inhibits PPAT enzymatic function is consistent with earlier reports that found reduced catalytic turnover with PPAT multimers (*43, 44*). In addition to direct inhibition of PPAT activity, such higher order structures might also prevent channeling of product to the next enzyme, GART, in purine *de novo* synthesis or interfere with purinosome formation. Regarding the molecular purine sensing mechanism, we find that the availability of salvageable adenosine increases the binding of NUDT5 to PPAT via the action of its PPAT interaction loop (Fig. 5f). We observe that NUDT5^Y74E^, but not NUDT5^Y74F^ abolishes PPAT binding, suggesting that the regulatory switch might occur by allosteric rearrangements or posttranslational modification of this residue (Fig. 5b and fig. S8c). Tyrosine phosphorylation of NUDT5^Y74^ has been reported in various proteomic studies (*45*), and future experiments will explore the impact of this modification.

Our findings led us to reevaluate the current model of *de novo* purine synthesis regulation. The current model suggests that PPAT is regulated via feedback inhibition of AMP and GMP at high µM to mM concentrations (*44*). However, we note that in our reporter assay conditions, MTHFD1^K56R^ cells are already sensitized to adenosine concentration in the low µM range (Fig. 1c), indicating a cellular purine sensing and NUDT5/PPAT dependent purine *de novo* biosynthesis regulation mechanism, that is several orders magnitude more sensitive than what the biochemical data obtained with PPAT alone suggest.

Furthermore, our findings can have translational applications. MTHFD1 deficiency is a rare genetic disease, and consistent with the presence of MTHFD1 C/H domain mutations we find adenosine to exert toxicity in patient cells. Possibly such effects can contribute to the observed immunodeficiencies as also other mutations of adenosine metabolism have been associated with such diseases.

Interestingly, NUDT5 has been identified to modulate the toxicity of the approved anticancer drug 6-TG. Here, we find that the rescue mechanism occurs through loss of PPAT binding rather than loss of the catalytic activity of NUDT5 itself (fig. S8d). These data suggest that modulation of purine *de novo* synthesis likely controls efficiency of purine analogs in anticancer therapy. This may have important clinical implications as drug resistance may evolve through loss of enzymatic HPRT1 function and/or emerging PPAT binding-deficient NUDT5 mutations.

Finally, with dNUDT5 we have generated a tool compound that enables pharmacologic modulation of the pathway, and future experiments will clarify whether such degraders will have translational applications to protect healthy tissues from chemotherapy side effects or MTHFD1deficiency patients from adenosine-mediated toxicities.

## Acknowledgments

We thank the Biomedical Sequencing Facility, the Proteomics and Metabolomics and Chemical Screening Facilities of the Molecular Discovery Platform at CeMM as well as I. Vendrell, S. Hester, R. Fischer and B. Kessler and the Proteomics Facility at the TDI in Oxford for their support in generating and analyzing the next generation sequencing or proteomics data, respectively. We are grateful to Jan-Lennart Venne for technical assistance and all members of the Kubicek and Huber labs for valuable discussions about the project.

## Author contributions

Conceptualization: JGL, TAN, SS, KVMH and SK

Methodology: JGL, TAN, ASMCM, YL, JH, LGB, LC, TJH, KK, KVMH, SK

Investigation: JGL, TAN, ASMCM, LDLD, YL, JH, LGB, JS, CB, LC, MS, JWG, GH, AK, KR, BG

Formal analysis: JGL, TAN, ASMCM, YL, JH, LGB, CB, LC, PB, MS, JWG, JTH, KK

Visualization: JGL, PB, LGB Resources: DSF, DSR

Funding acquisition: KVMH, SK

Supervision: TJH, KK, AB, JM, SS, KVMH, SK

Writing – original draft: JGL, TAN, ASMCM, LGB, KVMH, SK Writing – review & editing: all coauthors

## Data and materials availability

All cell lines generated in this study will be made available under an MTA. RNA-seq data have been deposited in GEO (accession number GSE201334).

## Materials and Methods

### Cell culture and transfection

HAP1 (KBM7-derived, Horizon Discovery, C859) cells and genetically modified variants (KO and lentiviral transduced cell lines) were cultured in Iscove’s Modified Dulbecco’s Medium (IMDM, Sigma), supplemented with 10% Fetal Bovine Serum (FBS, Sigma) and 1% Penicillin/Streptomycin (P/S, Sigma). Patient-derived fibroblast(*46*) and control fibroblast lines were cultured in Minimum Essential Medium alpha media (MEMα, no nucleosides, Gibco), supplemented with 10% (v/v) FBS and 1% (v/v) P/S. These patient-derived fibroblast lines were previously characterized. Dialyzed FBS (Gibco) was added to IMDM with 1% (v/v) P/S at a final concentration of 10% (v/v) for indicated experiments (DIA). HeLa and A549 cells were cultured in Dulbecco’s modified Eagle’s medium (DMEM, Sigma) supplemented with 10% FBS (Sigma) and 1% P/S (Sigma). All cell lines were incubated in 5% CO_2_ atmosphere at 37°C.

HAP1 cell lines were transfected with Turbofectin 8.0 (Origene) or Polyjet^TM^ (SignaGen Laboratories) according to manufacturer’s instructions.

### Plasmids and cloning

#### Cloning of MTHFD1 mutants

MTHFD1 cDNA was ordered from Genscript and cloned into lentiviral compatible vector (a gift from Eric Campeau & Paul Kaufman, Addgene plasmid # 17448) resulting in a N-terminal GFP-tagged construct. MTHFD1^K56R^ and MTHFD1^K386E^ mutants were created using the Q5 site-directed mutagenesis kit according to manufacturer’s instructions. For the genome-wide screen, the K56R mutant was also cloned into the lentiviral backbone with neomycin resistance (Addgene plasmid #17447, kind gift from Eric Campeau and Paul Kaufman).

#### Cloning of Intron tagging plasmids

The intron tagging plasmids were cloned as follows: The generic sgRNA targeting plasmid and the GFP-donor or mCherry-donor plasmid were generated as previously described (*47*). Briefly, the pX330 plasmid expression Cas9 and the generic sgRNA targeting the donor plasmid was generated by digesting pU6-(BbsI) CBh-Cas9-T2A-mCherry (Addgene #64324, kind gift from Ralf Kuehn) with BbsI followed by ligation with an annealed oligo duplex. mCherry was replaced with a Blasticidin resistance (BSD) using Gibson Assembly. The GFP-donor plasmid containing the coding sequence of EGFP flanked by generic sgRNA targeting sites, splice acceptor and slice donor sites and 20 amino acid linkers was assembled from 4 fragments using Gibson assembly. The DNA fragment with a 25 nucleotide overlap to the pUC19 vector and 32 nucleotides overlap to the N-terminus of EGFP was generated from overlapping oligos (Sigma) and is comprised of a generic sgRNA targeting site that is not present in the human genome followed by a splice acceptor site and a flexible 20 amino acid glycine-serine linker. This fragment is followed by a fragment with the coding sequence of EGFP without a start or stop codon that was generated by PCR. The third fragment has a 27 nucleotide overlap to the C-terminus of EGFP and a 25 nucleotide overlap to the pUC19 vector and was generated from overlapping oligos (Sigma) and comprises a flexible 20 amino acid glycine-serine linker followed by a splice donor site and the generic sgRNA targeting site. The pUC19 vector was linearized by PCR for Gibson Assembly (NEBuilder HiFi DNA Assembly) with the other three fragments. The mCherry-donor plasmid was similarly constructed.

#### Cloning of NUDT5 mutants

NUDT5^wt^ cDNA was obtained via RT-PCR (NEB, E3006) from isolated HAP1 RNA (Qiagen RNeasy kit). NUDT5^wt^ was amplified with primers harboring BsrGI_P2A (fwd) and SalI (rev) restriction sites and was cloned into BsrGI/SalI digested pLenti CMV_mcherry Puro vector. This construct was obtained from amplifying generic mcherry sequence with primers containing an XbaI (fwd) and SalI (rev) restriction site, which was cloned into pLenti CMV GFP Puro (Addgene plasmid # 17448). Respective point mutations were then cloned via Q5 Site-directed Mutagenesis Kit (NEB, E0554S) or site-directed ligation independent mutagenesis (*48*). HA_2_-NUDT5 was amplified from the mcherry_P2A_NUDT5 constructs by amplifying the NUDT5 sequence with primers containing an XbaI_HA_2_ (fwd) and SalI (rev) restriction site, which was then cloned into XbaI/SalI digested pLenti CMV_mcherry Puro backbone.

#### Cloning of plasmids for KO cell line generation

sgRNAs were designed using the online resource tool provided by Benchling (https://benchling.com, 2023) and purchased as short single-stranded DNA oligos with BbsI sticky ends. Oligonucleotides were phosphorylated, annealed and cloned into BBsI digested pX330 (mcherry selection, a gift from Ralf Kuehn, Addgene plasmid # 64324) or pX459 (puromycin selection, a gift from Feng Zhang, Addgene plasmid # 62988) backbone according to published procedures (https://portals.broadinstitute.org/gpp/public/).

### Generation of CRISPR–Cas9 genome edited cell lines

#### KO cell lines

WT or MTHFD1^K56R^ cells were transiently transfected with pX330/pX459 plasmid harbouring respective sgRNA sequence for protein target. Media was exchanges 24 h after transfection followed by 24 h of recovery. Cells were then selected either by sorting for expression of the respective fluorescence marker (Sony SH800S cell sorter) or antibiotic selection (2 µg/ml puromycin, 48 h). Cell pools were diluted single cell stage, expanded and individual clones validated by sequencing and western blot.

#### Intron tagged cell lines

The three plasmids (intron targeting, donor targeting, GFP/mCherry donor) were transfected into WT, MTHFD1^KO^, or MTHFD1^K56R^ cells. After 48 h, cells were collected and sorted by flow cytometry (Sony SH800S cell sorter). Cells were then diluted and split into single clones. After expansion, the colonies were visualized on an Opera Phenix (Revvity) to determine clones successfully tagged with the GFP/mCherry at the intron.

### Lentiviral transduction

Lentivirus was produced by co-transfecting HEK293T cells with pMD2.G (a gift from Didier Trono, Addgene plasmid #12259), psPAX2 (gift from Didier Trono, Addgene plasmid #12260) and lentiviral plasmid with PEI. 24 h after transfection, media was exchange to full IMDM followed by another 48 h of incubation. Lentiviral supernatant was then harvested, filtered (0.45 µm), aliquoted and flash frozen and stored at -80°C until further use. Lentivirus supplemented with 5 µg/ml (genome-wide KO screen) or 8 µg/ml (single mutant cell line generation) were transduced to respective cells for 24 h, followed by media exchange, recovery and selection or downstream experiments. Cells were selected either by antibiotic treatment (puromycin 1µg/ml) or sorting for the expression of fluorescence marker (Sony SH800S cell sorter). To determine MOI, lentivirus was titrated at different ratio to cells followed by selection. Fraction of positive cells was plotted against the respective lentivirus concentration. Transduced cells were analysed by sequencing and western blot.

### Genome-wide CRISPR Screens

The genome-wide CRISPR screen was performed as described in ref. (*49*). Briefly, MTHFD1^K56R^ cells were transduced with the virus harboring the Brunello library in media containing 5µg/ml Polybrene aiming for MOI of ∼1 and >500x gRNA coverage. After 48 h, the media was replaced with fresh media supplemented with 1µg/ml Puromycin. After another 48 h, cells were either collected (control condition) or grown in the following screening conditions: 1) DIA or 2) ADE (DIA with 50µM adenosine). Respective media was replaced every 3-4 days for a total of 14 days. Cells were then collected, and the genomic DNA was extracted. (DNeasy Blood and Tissue Kit, Qiagen). Guide RNA sequences were amplified and attached with adapters using primers adapted from ref. (*50*). Product was gel purified and sequenced (CeMM, Biomedical Sequencing Facility). Sequenced guides were analyzed with PinAPL-Py online tool (*51*).

### Western blot analysis

Unless otherwise specified, cells were lysed in RIPA buffer (50 mM Tris HCl, pH 7.4, 150 mM NaCl, 1% NP40, 0.25% Sodium deoxycholate, 1 mM EDTA, 0.1% SDS) supplemented with 1x protease inhibitor cocktail (Roche, cOmplete^™^, Mini, EDTA-free Protease Inhibitor Cocktail). Cells were lysed on ice for 30’ and then cleared by centrifugation. Protein amounts were determined by BCA assay (ThermoFisher, A32957) prior loading and separation by SDS-PAGE and transfer to PVDF membranes. Membranes were blocked in 5% milk-TBST (1% v/v) for 1 h at r.t. prior incubation with primary antibody at 4°C, o.n. Membranes were then washed three times with TBST and then incubated with HRP-conjugated secondary antibodies, diluted 1:10000 in 0.5% milk-TBST for at least 1 h at r.t. After washing with TBST (3x), membranes were developed using Clarity (Max)^TM^ Western ECL substrate (Bio-Rad) and imaged with ChemiDoc^TM^ MP imaging system (Bio-Rad). Western blots were stripped with mild stripping buffer (3x; 1.5% glycine (w/v), 0.1% SDS (w/v), 1% Tween-20 (v/v), pH 2.2), followed by washing, blocking and reprobing with target primary and secondary antibody.

The following antibodies were used: rabbit mAb HA-tag (1:5000, Cell Signaling Technologies (C29F4), #3724), rabbit mAb NUDT5 (1:2000, abclonal, A0609), rabbit pAb PPAT (1:2000, abclonal, A6698), mouse mAb GAPDH (1:5000, scbt (6C5), sc-32233), mouse mAb alpha-Tubulin (1:1000, Abcam, ab7291), mouse anti-rabbit IgG-HRP (1:5000, scbt, sc-2357), goat anti-mouse IgG-HRP (1:5000, invitrogen, 31430).

### High throughput compound screen

MTHFD1^KO^ cells (F5 clone) in dialyzed FBS media were treated with the CeMM inhouse compound library (89,228 chemically diverse compounds). Both cell number, using Hoechst staining, and CellTiter-Glo (Promega), an indicator of metabolically active cells, was both used as a readout, depending on the stage of the screen.

Briefly, the screening was divided into the three parts (i) primary screening, (ii) follow-up and (iii) validation. During the primary screen, MTHFD1^KO^ cells were treated with 10 µM of every compound and CellTiter-Glo (Promega) was used to assess their ability to increase cell growth after 48 h. From this primary screening, 17 compounds were selected as hits and rescreened in the follow-up part, in which the MTHFD1^KO^ cells were treated in a six-point dose response for 72 h to discard any false positives. Finally, in the validation part, MTHFD1^KO^ cells were treated with 10 compounds in an eight-point dose response curve and the cell numbers were counted by staining with Hoechst, imaging and counting nuclei using the Opera Phenix (Perkin Elmer) and Harmony software (Perkin Elmer).

Compounds were transferred on 384-well plates using an acoustic liquid handler (Echo 550, Labcyte/Beckman Coulter) and 1,000 cells per well were dispensed on top of the drugs using a dispenser (Multidrop, Thermo Fisher Scientific) for a total of 50 μl/well. ATP levels was measured after 48 h using CellTiter-Glo (Promega) in a multilabel plate reader (EnVision, Perkin Elmer). Signal was then normalized to DMSO and adenosine control wells included on each plate.

### Single metabolite supplementation screen

Nucleotide metabolites (adenine, adenosine, guanine, guanosine, cytidine, cytosine, uridine, uracil, thymine, 5-methyluridine) were purchased from Sigma and dissolved in DMSO at a concentration of 50 mM. Deoxynucleotides (dATP, dCTP, dTTP, dGTP) in 100 mM solutions were purchased from Sigma. Folate metabolites (folic acid from Sigma, 5-formyl tetrahydrofolic acid, 5-methyltetrahydrofolic acid, 5,10-methenyl tetrahydrofolic acid, 5,10-methylene tetrahydrofolic acid, tetrahydrofolic acid, and dihydrofolic acid purchased from Schircks Laboratories) were dissolved in DMSO at a concentration of 40 mM.

MTHFD1^KO^ cells in dialyzed FBS media were treated with a single nucleotide or folic acid metabolite for 72 hours (50 µM). Cells were then stained with Hoechst, imaged, and counted using an Operetta (Perkin Elmer). Similarly, MTHFD1^K56R^ cells in dialyzed FBS media were treated with adenosine and nucleoside analoga (50 µM) for 72 h. Cells were then stained with Hoechst, imaged, and counted using an Operetta (Perkin Elmer).

### RNA-seq and GO Term Enrichment Analysis

Patient cells were incubated in FULL, DIA, or ADE media for 24 h and then the RNA was extracted using the RNeasy Mini kit (Qiagen). RNA was sequenced by the Biomedical Sequencing Facility at CeMM using the Illumina HiSeq3000/4000 platform and the 50-bp single-end configuration. Reads were aligned with TopHat (*52*). Cufflinks (*53*) was used to assemble aligned RNA-Seq reads into transcripts, estimate their abundances, and test for significant differential expression. Genes were significant if the calculated q-value was less than 0.05. RNAseq data is stored with GEO study accession number GSE201334.

For the GO term enrichment analysis, up-regulated and down-regulated proteins were defined based on log(FC) for each experimental condition. The enrichment analysis for resulting list of proteins was performed using enrichr API (https://maayanlab.cloud/Enrichr/) (*54, 55*) through ‘enricher’ package in R (https://cran.r-project.org/web/packages/enrichR/index.html) with “GO_Biological_Process_2018” library.

### Immunofluorescence staining

Cells were pretreated in different media conditions for 24 hours and then fixed in 4% paraformaldehyde, and permeabilized with Triton-X. The fixed cells were then blocked with 5% BSA, diluted in PBS, for 1 hour and then incubated overnight with anti-γH2A.X antibody (1:500, 9718S, Cell Signalling Technology) or rabbit anti-RPA2 antibody (1:500, HPA026306, Atlas Antibodies) at 4°C. The next day, the cells were washed and incubated with anti-rabbit Alex-Fluor 546 (1:1000, A11010, Thermo Fisher Scientific) or anti-rabbit Alexa Fluor 568 (1:1000, A11036, Thermo Fisher Scientific) and Hoechst 33342 (1 µg/ml, HY-15559A, MedChem Express) for 1 h at r.t. The cells were washed again with PBS prior to imaging. Foci were imaged using Opera Phenix (Revvity) and quantified using Harmony software (Revvity) or a custom CellProfiler pipeline (*56*).

### Cell cycle analysis

Cells were pretreated in indicated media conditions for 24 hours, trypsinized and washed twice with ice-cold PBS. After washing, cells were resuspended in 0.5 ml of ice-cold PBS. 2 ml of ice-cold 70% EtOH was then added dropwise to the cells, while gently vortexing. Cells were then stored at -20°C for at least 20 minutes. After fixation, cells were resuspended in PBS for 5 minutes and then stained with propidium iodide (Cell Signaling Technology, #4087S) according to manufacturer’s protocol. DNA content was measured with Sony SH800S cell sorter and analyzed with FlowJo^TM^.

### Targeted metabolomics and stable isotope tracing

All cell lines were incubated in the respective media conditions: full FBS media, dialyzed FBS media, and dialyzed FBS media supplemented with 1 mM isotope labelled sodium formate (Cambridge Isotope Laboratories, CDLM-6203-0.5) or 1 mM isotope labelled sodium formate and 50 µM isotope labelled adenosine (Silantes,125303601) for 24 h prior to collection. Cells were counted and 5 million cells for each replicate were washed with 1x PBS, pelleted, immediately snap-frozen, and stored at -80°C.

Cell extraction was performed by adding 500 µL of ice-cold 80/20 MeOH/H_2_O (v/v) solution to the cell pellet while vigorously vortexing. Samples were centrifuged at 10,000 g for 10 min at 4°C before transferring the cell extract supernatant into 1.5 ml HPLC vials. The extraction of the cell pellets was repeated a second time and supernatants of the same samples were combined.

Cell extract samples were dried using a nitrogen evaporator. The dried residue was reconstituted in 50 µL water. An aliquot of 10 µL reconstituted sample extract was mixed with 10 µl of isotopically labelled internal standard mixture in a HPLC vial, vortexed and used for the metabolite analysis.

A 1290 Infinity II UHPLC system (Agilent Technologies) coupled with a 6470 triple quadrupole mass spectrometer (Agilent Technologies) was used for the LC-MS/MS analysis. The chromatographic separation for samples was carried out on a ZORBAX RRHD Extend-C18, 2.1 x 150 mm, 1.8 µm analytical column (Agilent Technologies). The column was maintained at a temperature of 40 °C and 4 µL of sample was injected per run. The mobile phase A was 3% methanol (v/v), 10 mM tributylamine, 15 mM acetic acid in water and mobile phase B was 10 mM tributylamine, 15 mM acetic acid in methanol. The gradient elution with a flow rate of 0.25 ml/min was performed for a total time of 24 min. Afterwards a back flushing of the column using a 6-port/2-position divert valve was carried out for 8 min using acetonitrile, followed by 8 min of column equilibration with 100% mobile phase A.

The triple quadrupole mass spectrometer was operated in an electrospray ionization negative mode, spray voltage 2 kV, gas temperature 150 °C, gas flow 1.3 l/min, nebulizer 45 psi, sheath gas temperature 325 °C, sheath gas flow 12 l/min. The metabolites of interest were detected using a dynamic MRM mode. The MassHunter 10.0 software (Agilent Technologies) and PeakBotMRM (https://github.com/christophuv/PeakBotMRM) were used for data processing and stable isotope tracking. Ten-point linear calibration curves with internal standardization were constructed for the quantification of metabolites.

### Procedures for synthesis of dNUDT5

The synthesis and detailed characterization of dNUDT5 is described in the accompanying manuscript *Marques et. al.* submitted to Nature Chemical Biology.

### Immunoprecipitation

#### Co-immunoprecipitation

MTHFD1^K56R^ cells were cultured in 10 cm dishes in DIA and transfected with HA_2_-NUDT5 constructs. After 24 h, media was exchanged, and cells were grown for another 24 h prior harvesting. For adenosine dependent binding studies, cells were additionally treated with 50 µM adenosine for 3 h. Cells were washed twice with ice-cold PBS, evenly divided in three aliquots, pelleted, flash frozen and stored at -80°C until further use. One aliquot of cell pellets was lysed in TBST (1%) with 0.05% NP40, 1x protease and phosphatase inhibitors (ThermoFisher scientific, A32963) for 30’ at 4°C followed by clearing via centrifugation. Equal amounts of lysates were incubated with 2x washed (20 mM Tris HCl, 150 mM NaCl, pH 7.4 - washing buffer) anti-HA magnetic beads (Sigma-Aldrich, SAE0197) for 1 h at 4°C while being rotated. Beads were washed twice with washing buffer and once with water prior elution in Laemmli buffer at 95°C for 10’. SDS-PAGE and western blot were performed and developed as described above.

#### BFP-pulldown

MTHFD1^K56R^NUDT5^KO^ cells and cells reconstituted with BFP-NUDT5 (10^7^ cells) were lysed and processed as described above. 20 µl of GFP selector resin (NanoTag Biotechnologies, N0510-L) slurry was washed with twice with lysis buffer according to manufacturer’s protocol and 2.5 mg of lysate was added to the beads. After 1 h incubation at 4°C while being rotated, beads transferred to a Mini Spin column (BioRad, 7326204) and washed twice with wash buffer (lysis buffer without protease inhibitor) and once with Tris-HCl (100 mM, pH 7.8) according to manufacturer’s protocol. Beads were incubated with lysis buffer supplemented with 4% SDS at 95°C for 2 min. Eluates were captured and subjected to proteomic analysis (CeMM Molecular Discovery Platform).

#### NUDT5 TurboID

HEK293 FlipIn cells were transfected with TurboID-NUDT5 and pOG44 using Lipofectamin2000. After 5 days, cells were selected using blasticidin (100 µg/mL) and hygromycin (10 µg/mL) for ca. 2 weeks. Cells were seeded in 10 cm dishes and incubated overnight. The next day 1.3 µg/ml doxycycline was added, incubated overnight, before changing the medium to DMEM + 10% dialysed FBS, 1.3 µg/ml doxycycline, and 50 mM adenosine. After 24 h incubation, 50 mM biotin was added, incubated for 4 h, before harvesting. Pellets were washed 2x with PBS and then stored at -80 °C. Pellets were lysed with RIPA buffer containing 1 µg/µl benzonase for 30 min on ice. Lysates were cleared by centrifugation at top speed in a tabletop centrifuge at 4°C. Protein concentrations were measured using a DC assay and 3 mg per sample in 850 µl were mixed with 150 µl streptavidin beads and incubated on a wheel at 4 °C over night. Beads were washed with 3x 1 ml RIPA buffer, 1x 1 ml 1 N KCl, 3x 1 ml PBS, before resuspension in 50 µl 1x Laemmli. Beads were boiled at 95 °C for 10 min, eluates collected, and the elution procedure repeated one more time.

#### PPAT Co-Immunoprecipitation

WT cells were treated with 100 nM TH5427, dNUDT5 or DMSO for 20 h in a 100 mm dish. Cells were washed four times with 2 ml PBS before lysing with 500 µl 0.8% NP-40 based lysis buffer (Tris [pH 7.5], 0.8% NP-40, 5% glycerol, 1.5 mM MgCl_2_, 100 mM NaCl, 25 mM NaF, 1 mM Na_3_VO_4_, 1 mM PMSF, 1 mM DTT, 10 µg/ml TPCK, 1 µg/ml Leupeptin, 1 µg/ml Aprotinin, 10 µg/ml soybean trypsin inhibitor). The cell suspension was transferred to a 2 mL reaction tube, incubated on ice for 30 min and then centrifuged for 30 min at 20000 xg (4 °C). Cleared lysates were spiked with 1 µg of PPAT (15401-1-AP, Proteintech) or rabbit IgG control antibody (3900S, Cell Signaling) and incubated on a wheel at 4 °C overnight. 30 µl magnetic protein G beads slurry (#1003D, ThermoFisher) was incubated with lysate/antibody mix for 15 min on room temperature on a wheel. After 3x washing with 1 ml lysis buffer protein were eluted with 30 µl 2x Laemmli buffer by boiling for 10 min at 70 °C.

#### Chemical pulldown

Profiling of the TH5427 proteome wide specificity was performed in triplicates as previously described (*57*). In brief, NHS-activated sepharose beads (17090601, Cytiva) were derivatised with an amine functionalised TH5427 analog (CBH-003) (*58*) (manuscript in preparation) at a coupling density of 0.5 mM with 0.75 µL TEA per 50 µl beads in DMSO. Free binding sites were blocked by 2.5 µL ethanolamine per 50 µl beads. Beads were washed with excess of DMSO and 0.8% NP-40 based lysis buffer. WT cell pellets were thawed on ice and lysed with 3x excess lysis buffer containing 1 µl/1 ml benzonase. The cell suspension was drawn through a 21G needle 10x before clearing cell debris by centrifugation for 30 min at 20000 × g (4 °C) in a table-top centrifuge. 20 µM of TH5427 or DMSO was spiked into 1 ml of lysate (5 mg/ml) and incubated for 30 min on a wheel at 4 °C. After mixing cell lysates with 50 µl beads for 2 h on a wheel at 4 °C the suspension was transferred in filter columns, washed 4x 1 ml lysis buffer and subsequently eluted with 2x 50 µl 2x Laemmli buffer by heating for 5 min at 95 °C. Elution fractions were used for gel loading or digested for LC-MS/MS analysis.

### Proteomics

#### Untargeted global proteomics and chemoproteomics

Proteomics samples were digested with trypsin using STrap columns following the manufacturer’s protocol (C02-micro, ProtiFi). Approximately 80% of the elution fraction of the pulldown were spiked with 5% SDS, then reduced by 20 mM dithiothreitol (M02712, Fluorochem), and alkylated by 40 mM iodoacetamide (I1149, Sigma). Phosphoric acid and S-Trap protein binding buffer were added into the sample lysates. Then, the SDS lysate/S-Trap buffer were loaded on a S-Trap column (C02-micro-80, ProtiFi). 1 μg of trypsin (V5111, Promega) was used to digest each sample at 37 °C for 20 h. The samples were then eluted and dried by vacuum centrifugation. Peptide pellets were resuspended in mass spectrometry grade water with 2 % acetonitrile (85188, Thermo Scientific) and 0.1 % trifluoroacetic acid (85183, Thermo Scientific). The samples were injected into Orbitrap Fusion Lumos Tribrid Mass Spectrometer (Thermo Scientific). Sample acquisition was performed as previously described (*59*).

#### Data analysis

Raw data were searched against the human database (UP000005640, downloaded 08/21) using DIA-NN (v.1.8.1) (*60*) by enabling FASTA digest for library free search and Deep learning-based spectra, RTs and IMs prediction. Trypsin/P was selected for the in-silico digestion of the provided fasta file, allowing up to 2 missed cleavages. Fixed modifications included N-terminal methionine excision and cysteine carbamidomethylation. Variable modifications were methionine oxidation and N-terminal acetylation, with a maximum of one modification per precursor. Precursor FDR was kept at 1% and MBR enabled. The protein groups matrix was further analysed using Perseus (2.0.9.0) (*61*) or R studio (4.3.2). Potential contaminants were filtered against the top 300 most common Homo sapiens contaminants reported in CRAPome (*62*). Proteins were removed if detected less than 50% across all runs. After column-wise normalisation by median subtraction, missing values were imputed by random numbers from a normal distribution (1.8 standard deviation, downshift width 0.3). Volcano plots were generated by calculating fold change and -log10(p-value) from a two-sided Student’s t-test.

### AlphaFold modelling

AlphaFold models were generated using AlphaFold3 and the default settings (*38*). Four units of NUDT5 (UniprotID: Q9UKK9) and PPAT (UniprotID: Q06203) respectively were submitted and resulting structures were analyzed with Pymol.

### Fluorescence competition assay

MTHFD1^K56R^ NUDT5^KO^ cells were transduced with lentiviral packed mcherry_P2A_NUDT5 constructs supplemented with 8µg/ml polybrene, aiming for an MOI <1. After 24 h, media was exchanged, and cells were expanded for another 24 h. Equal amounts of cells were then seeded in 6 well plates in DIA media. Next day, media was replaced with DIA or ADE media followed by cultivation for 72 h. Cells were harvested and subjected to FACS analysis (LSRFortessa^TM^, BD Biosciences). Ratio of mcherry positive cells was determined by normalizing the values of ADE treated cells to DIA control conditions.

### Cell viability assay

#### Adenosine toxicity

Cells were cultured in dialyzed media in 96 well plates (1200cells/100µl/well) supplemented with or without 50 µM adenosine. After 72 h incubation, media was exchanged to dialyzed with Hoechst 33343 (1 µg/ml) and after 10’ of incubation nuclei were imaged and counted (Opera Phenix, Revvity).

#### 6-thioguanine assay

Cells were cultured in full media in 96 well plates (1200cells/100µl/well) supplemented with or without indicated concentration of 6TG. After 72 h incubation cell viability was measured using CellTiter-Glo assay (Promega). Signals were normalized to control condition.

### Statistical Analysis

All statistical tests and plots were performed in Prism (Prism 10). For comparisons among three or more conditions, ANOVA was performed with Tukey’s HSD to correct for multiple comparisons. For comparisons between two conditions, T-tests were performed with a two-tailed p-value.

---

**Fig. S1.**
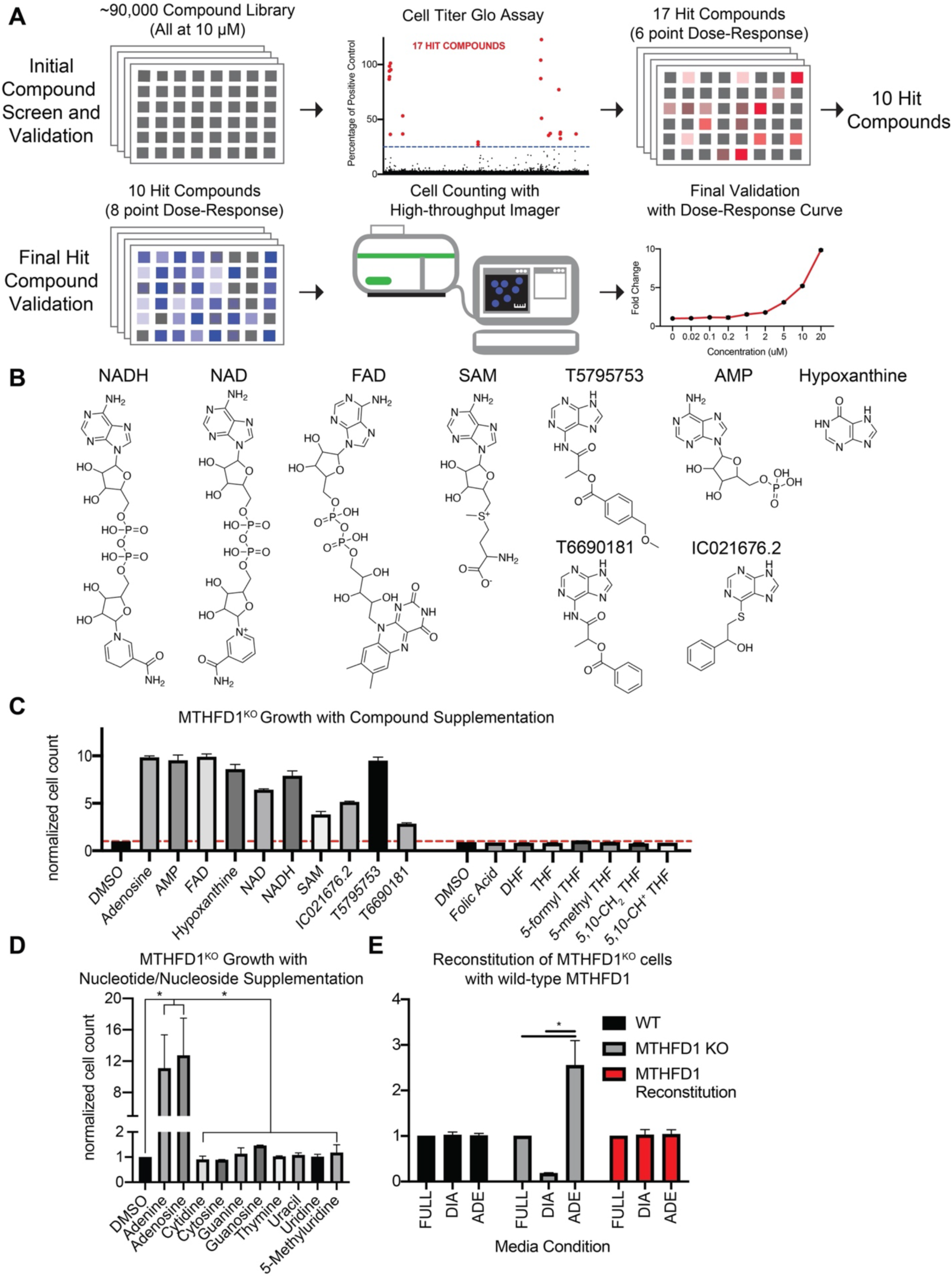
Large-scale compound screen reveals that only adenine-containing compounds rescue the growth of MTHFD1^KO^ cells. **(A)** Schematic of the two-stage compound screen workflow. (**B**) Chemical structures of the validated hit compounds. (**C)** Normalized cell numbers of MTHFD1^KO^ cells grown in dialyzed media supplemented with the validated hits from the screen and folate precursors (n=2 biological replicates, mean ± s.d., *p<0.05) **(D)** Normalized cell numbers of MTHFD1KO cells grown in dialyzed media supplemented with indicated nucleosides and nucleotides (n=2 biological replicates, mean ± s.d., *p<0.05). **(E)** MTHFD1 reintroduction into MTHFD1^KO^ cells abolishes effects of dialyzed serum and adenosine supplementation.

**Fig. S2.**
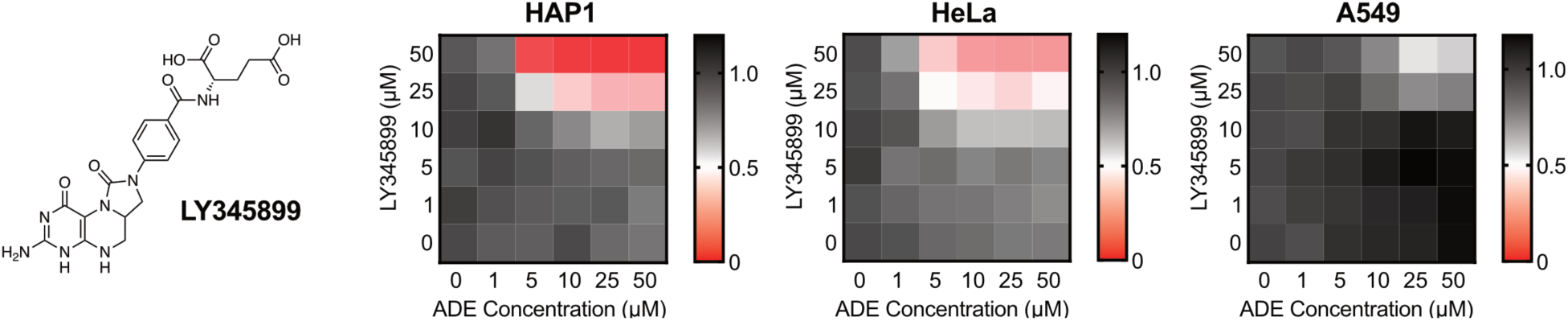
Inhibition of the MTHFD1 D/C domain sensitizes selected cancer cell lines to adenosine. Normalized cell counts of HAP1, HeLa and A549 cells treated with indicated concentrations of MTHFD1 D/C domain inhibitor LY345899 and adenosine for 72 h. Cell counts were normalized to the DMSO condition for each cell line. (n=2 biological replicates).

**Fig. S3.**
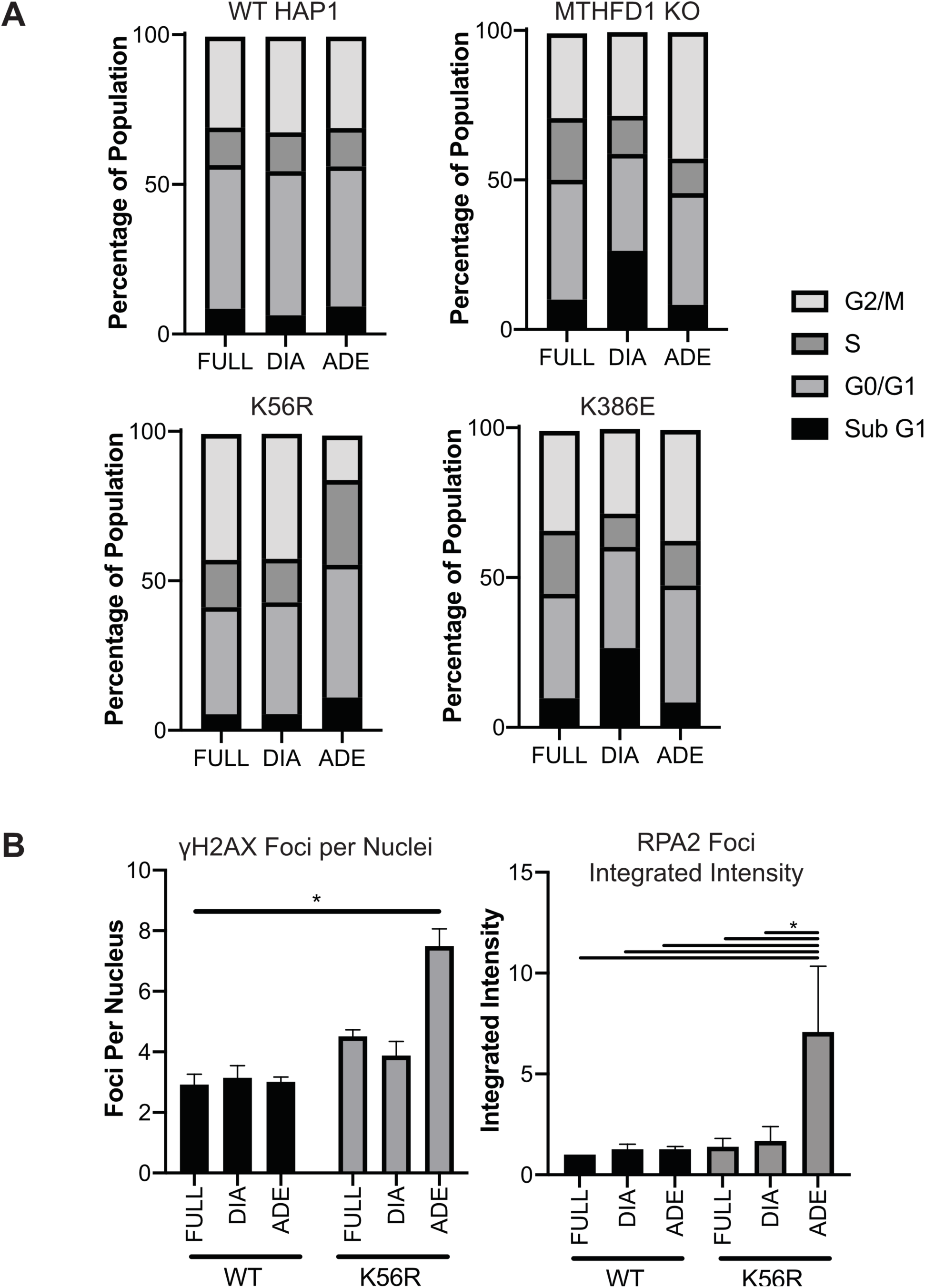
Quantification of cell cycle and DNA damage effects upon adenosine supplementation. **(A)** Quantification of the percentage of the depicted cell lines in specific cell cycle states. Cells were pretreated in FULL, DIA, or ADE for 24 h. (n=3 biological replicates, mean ± SD). **(B)** Quantification of the average number of γH2A.X nuclear foci and the integrated intensity (size and brightness) of RPA2 foci of HAP1^WT^ and MTHFD1^K56R^ cells grown in FULL, DIA, or ADE media (n=3 biological replicates, mean ± s.d., *p<0.05).

**Fig. S4.**
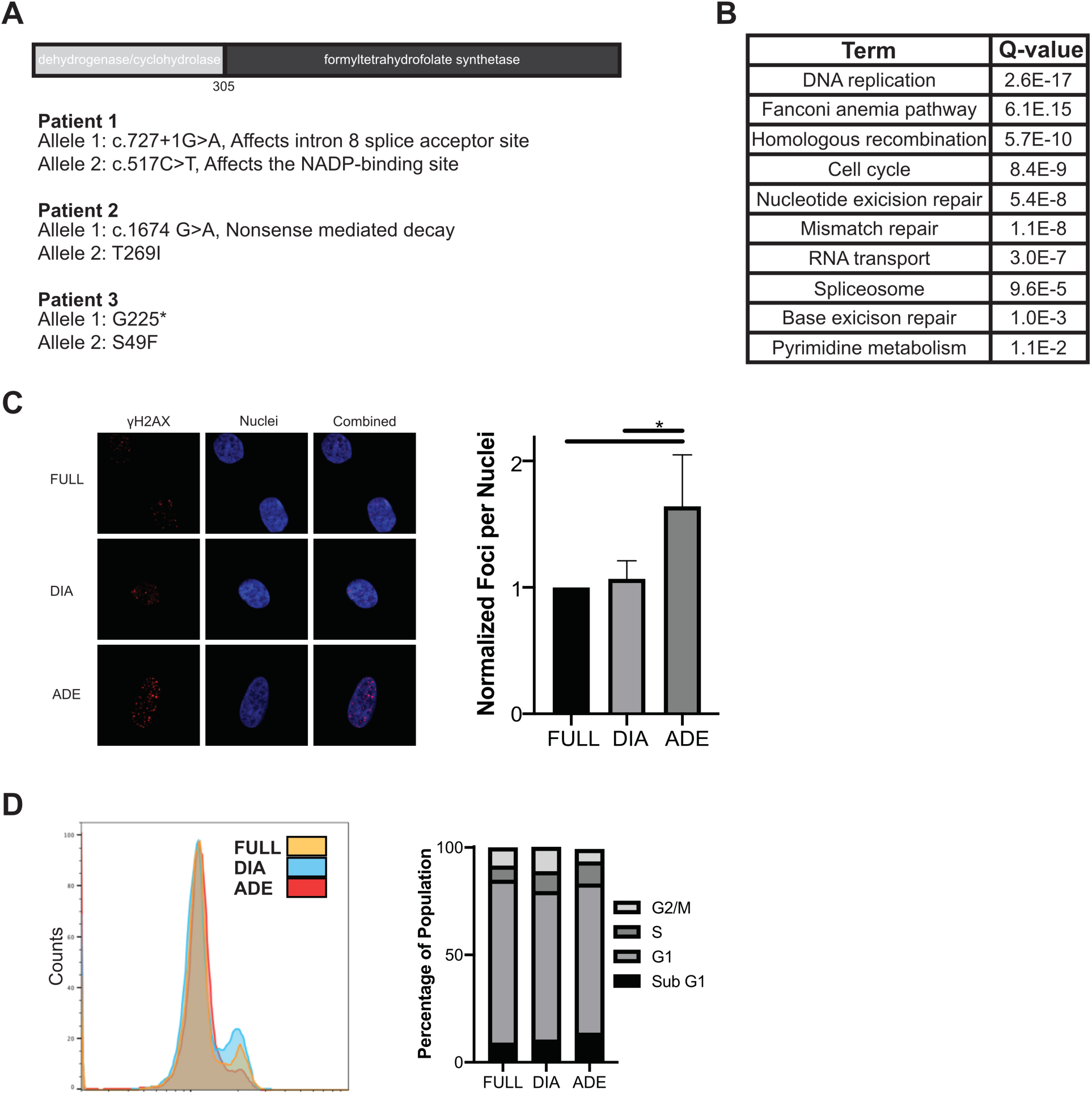
Adenosine supplementation leads to DNA damage and cell cycle arrest in MTHFD1-deficient, patient-derived fibroblasts. **(A)** Schematic representation of MTHFD1 domain architecture and characterized patient-specific mutations. **(B)** Highest enriched GO terms comparing the transcriptome of patient cells in ADE versus DIA media. **(C)** Representative images and quantification of the γH2A.X foci per nuclei of patient 2 cells pretreated in FULL, DIA, ADE media for 24 h. (n=3 biological replicates, mean ± SD, *p<0.05). (**D**) Cell cycle analysis of patient 2 cells pretreated in FULL, DIA, ADE media conditions. (n=3 biological replicates, mean ± s.d, *p<0.05)

**Fig. S5.**
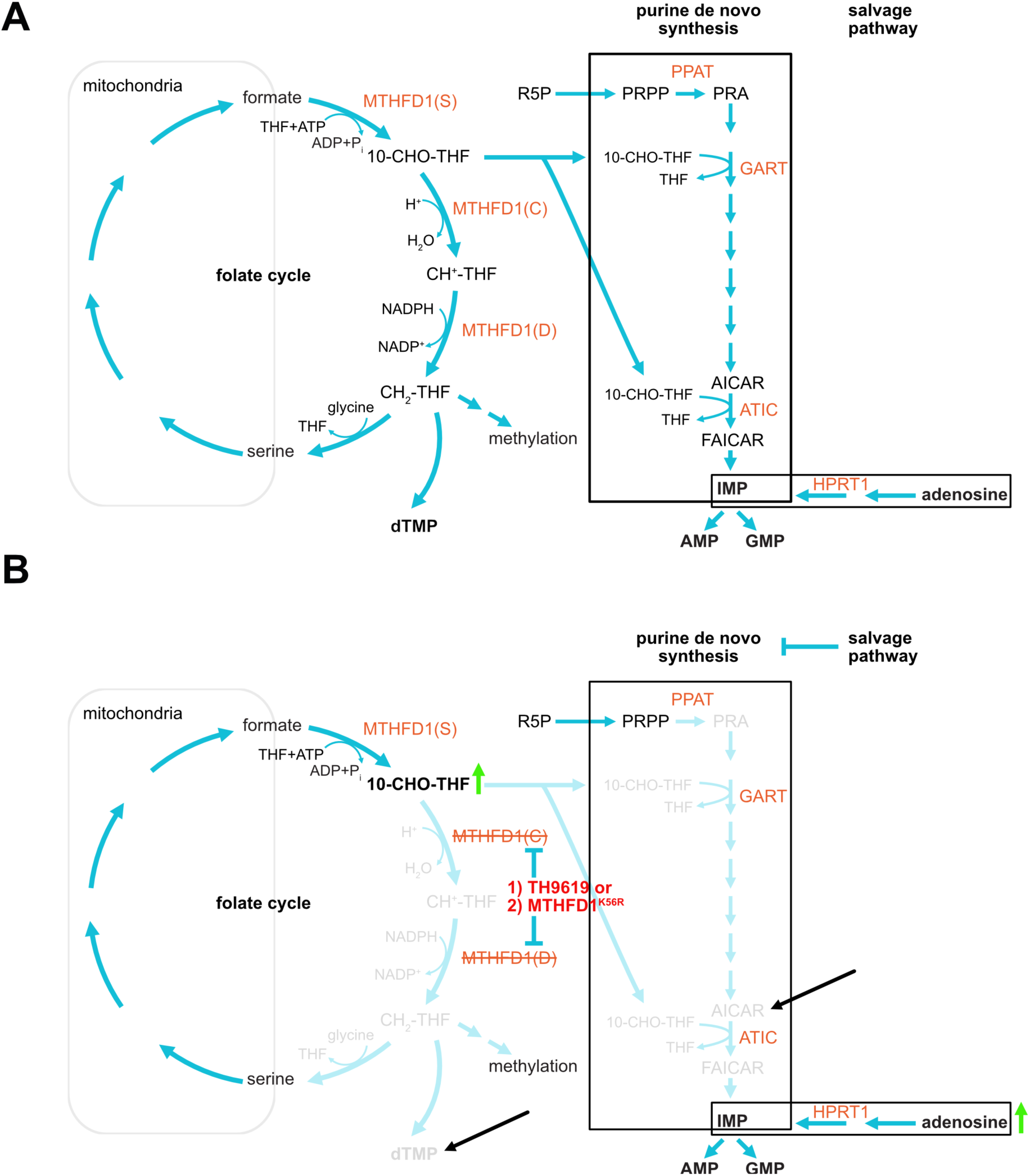
Detailed folate trap model. **(A)** Folate cycle, purine *de novo* synthesis pathway and salvage pathway are interconnected to enable cellular purine synthesis. **(B)** Proposed folate trap mechanism: Adenosine supplementation or elevated purine levels results in blocking of the purine *de novo* synthesis pathway. Simultaneous a) pharmacological (by TH9619) or b) genetic (by MTHFD1^K56R^ mutation) inhibition of MTHFD1 D/C domain causes accumulation of 10-CHO-THF and blocking of the folate recycling, thymidylate biosynthesis pathway. The resulting nucleotide imbalance is proposed to cause DNA damage and ultimately cell death. Black arrows indicate potential points of rescue by metabolite addition. AICAR supplementation and consequently conversion to FAICAR by ATIC requires 10-CHO-THF, which restores depleted THF levels while supplementation of thymidine (derivatives) presumably directly counteracts nucleotide imbalance and DNA damage. PRPP - Phosphoribosyl pyrophosphate, PRA – phosphoribosylamine, AICAR - 5-aminoimidazole-4-carboxamide ribonucleotide, FAICAR - 5-Formamidoimidazole-4-carboxamide ribonucleotide, GART - Phosphoribosylglycinamide synthetase, ATIC - 5-aminoimidazole-4-carboxamide ribonucleotide formyltransferase/IMP cyclohydrolase.

**Fig. S6.**
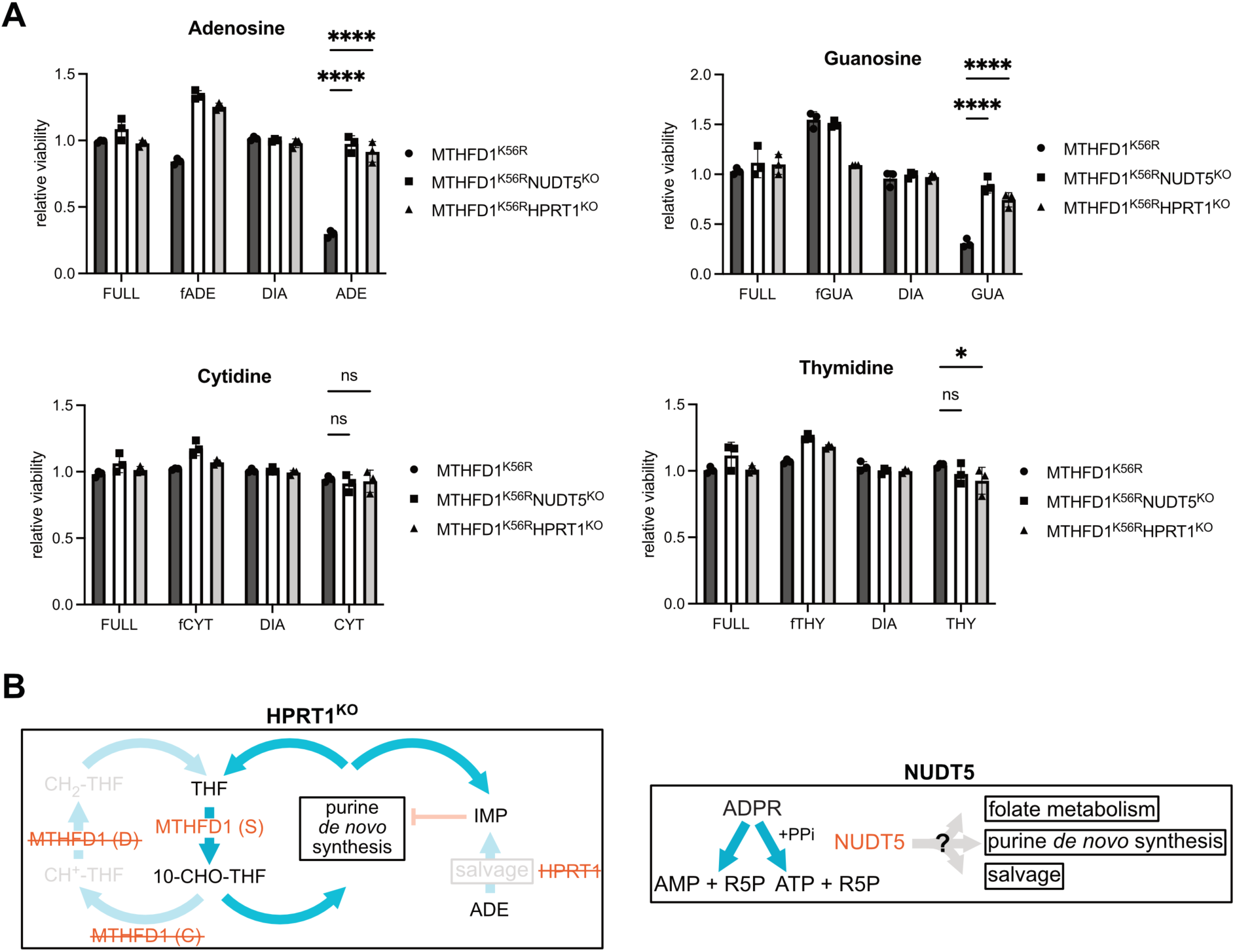
Nucleoside treatment of MTHFD1^K56R^ cells reveal purine over pyrimidine sensitivity. **(A)** MTHFD1^K56R^ cells were cultured in respective media and 50 µM nucleosides for 72 h prior CTG measurement (fADE – FULL supplemented with 50µM adenosine, similar for other nucleosides). Generally, the cells are sensitized to purine addition, while addition of pyrimidines showed toxicity effects. MTHFD1^K56R^HPRT1^KO^ and MTHFD1^K56R^NUDT5^KO^ reversed the purine mediated toxicity effects. (n=3, mean+s.d. ns – not significant, *p=0.0245 (Thymidine), ****p<0.0001) **(B)** Model for the proposed folate trap rescue in MTHFD1^K56R^HPRT1^KO^ cells and schematic overview of the enzymatic function of NUDT5 with putative connection to folate metabolism, purine *de novo* synthesis and salvage pathway (ADPR – ADP-ribose, R5P – ribose-5-phosphate).

**Fig. S7.**
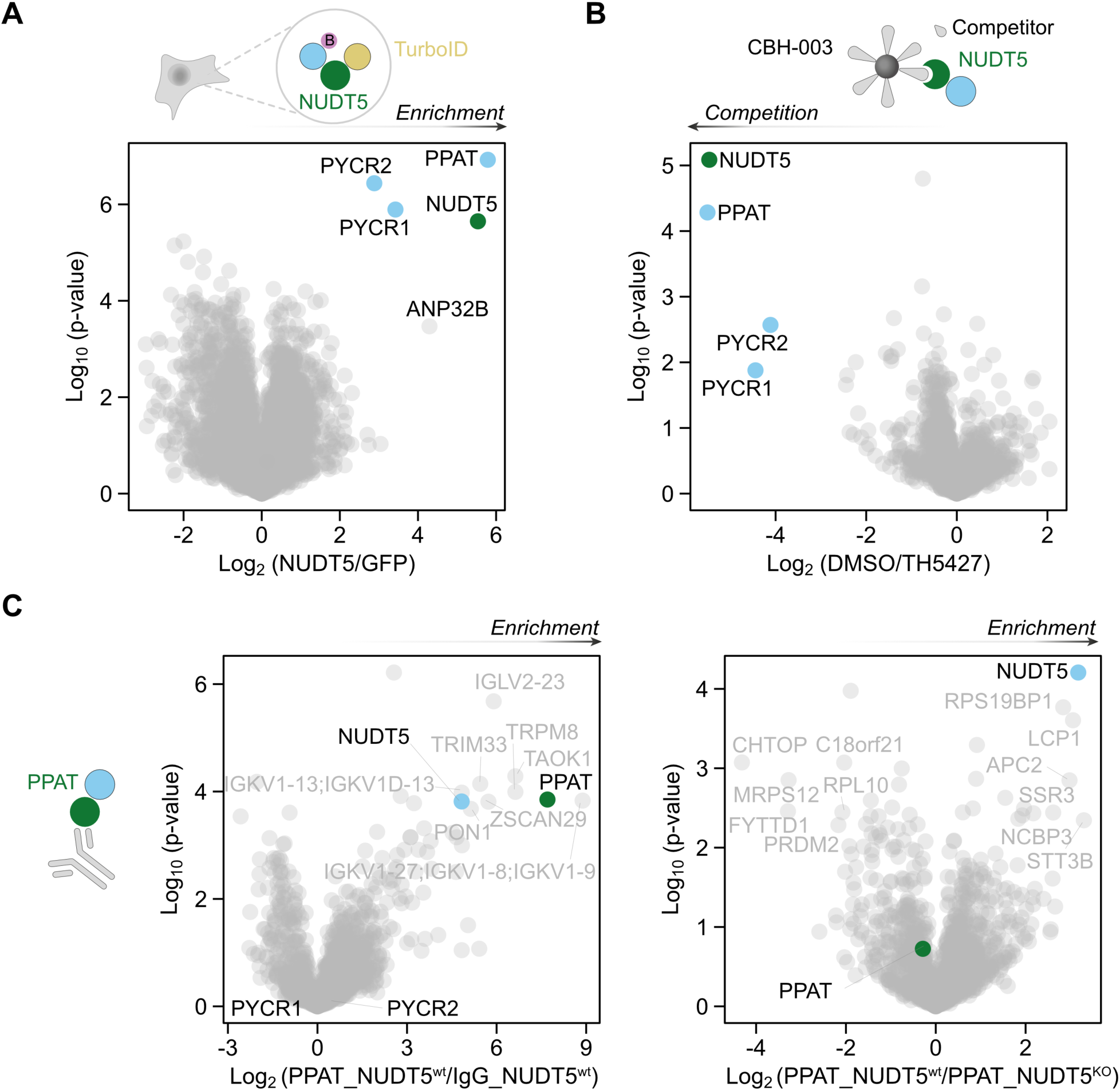
Large-scale interaction proteomic profiling of NUDT5 and PPAT interactors. The NUDT5/PPAT interaction was detected using various methods: **(A)** via NUDT5 TurboID proximity labelling in intact HEK293 FlipIn cells, **(B)** chemoproteomics using CBH-003 affinity matrix and TH5427 as competitor with HAP1 lysate and **(C)** PPAT Co-IP in HAP1 cells (left: PPAT vs. IgG in NUDT5^wt^, right: PPAT in NUDT5^wt^ versus PPAT in NUDT5^KO^)

**Fig. S8.**
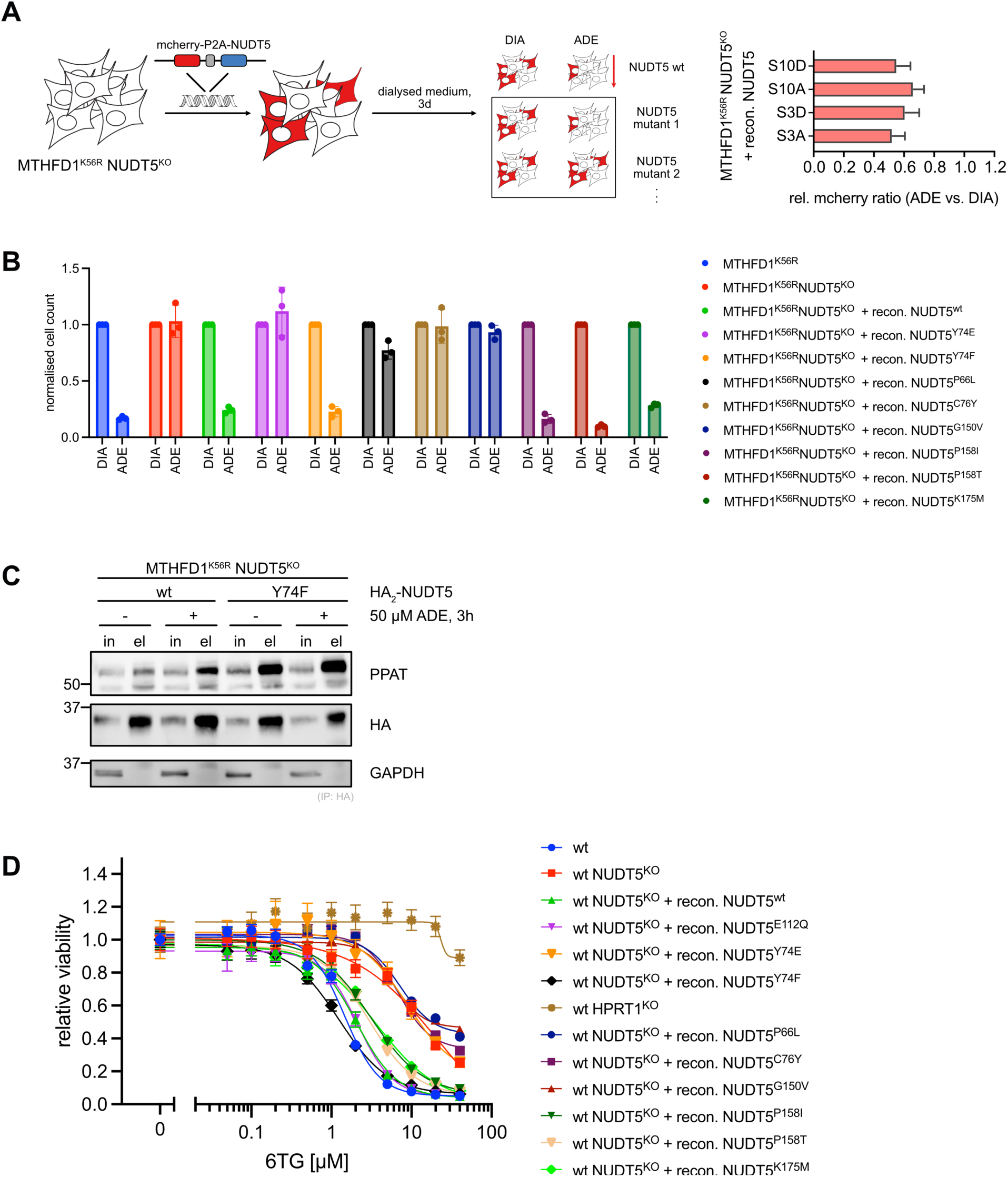
Selective NUDT5 mutations modulate PPAT interaction and purine and 6-TG sensitivity. **(A)** Schematic overview of fluorescence competition assay set-up to test the effects of various NUDT5 mutations on adenosine mediated toxicity in MTHFD1^K56R^ cells. Results of fluorescence competition assay with NUDT5 S3 and S10 (putative) phosphorylation inert (S→A) and mimic variants (S→D) are shown on the right. **(B)** Normalized cell counts of MTHFD1^K56R^, MTHFD1^K56R^NUDT5^KO^, and sorted pools of MTHFD1^K56R^NUDT5^KO^ cells reconstituted with indicated NUDT5 variants after being cultured in DIA and ADE for 72 h. **(C)** Pulldown of MTHFD1^K56R^NUDT5^KO^ cells transfected with HA_2_-NUDT5^wt^ or -NUDT5^Y74F^. Cells were cultured in DIA with or without adenosine (50 µM, 3 h; representative blot from two independent experiments). **(D)** Viability assays of HAP^wt^, NUDT5^KO^, HPRT1^KO^ cells and sorted pools of NUDT5^KO^ cells reconstituted with indicated NUDT5 variants at indicated concentrations of 6-TG.

